# Asymmetric Neurogenomic States Emerge in Winners and Losers After Social Competition

**DOI:** 10.64898/2026.07.26.740825

**Authors:** Ming-Tzu Chiu, Vu Trieu-Duc, Akiko Maruko, Kenshiro Oshima, Hao-Ven Wang, Norihiro Okada

## Abstract

Winner and loser effects can influence future aggressive behavior following social competition, but their neurogenomic basis remains poorly understood. Using *Betta splendens*, we integrated behavioral analyses, whole-brain RNA sequencing, and genomic coexpression network analyses to investigate how social experiences influence enduring behavioral and molecular states. Winners showed increased attack frequency, whereas losers prolonged aggressive latency, indicating effects on distinct behavioral aspects. Winners and losers initially shared acute responses in terms of both gene expression and network coordination but subsequently diverged through outcome-specific patterns of network reorganization, despite few persistent differentially expressed genes. This divergence was asymmetric and involved multiple components: immune-related networks showed notable contrasts, with winners displaying anticipated postconflict coordination but losers instead showing discoordination, whereas neuroendocrine and purinergic networks diverged over distinct temporal scales. These findings highlight transcriptomic network-level remodeling as a potential key feature of the winner–loser effect.

## INTRODUCTION

The outcome of social competition not only determines an animal’s social rank and access to resources but also shapes its performance in future conflicts [1,2]. Individuals with winning experience are more likely to win subsequent encounters, whereas those with losing experience often show reduced competitive performance. These winner–loser effects have been demonstrated across taxa ranging from insects to vertebrates and are considered a widespread form of experiment-dependent behavioral plasticity [1,3–7]. Because they influence social hierarchy formation, resource acquisition, and ultimately fitness, winner–loser effects have become an important model for understanding how social experience shapes behavior [1,8–10]. Despite extensive investigations at both the behavioral and physiological levels, important questions remain regarding both the behavioral organization and the molecular basis of these effects.

Winner and loser effects are usually treated as opposite ends of a single aggression continuum, with winners becoming more aggressive and losers less aggressive [1]. However, studies that assess multiple behavioral components reveal a more complex picture: the measures most responsive to winner effects differ from those most responsive to loser effects, and certain measures show an effect in one group but not the other [11,12]. In particular, loser effects on aggressive latency can persist even after the effects on attack frequency have diminished [1,11]. These observations suggest that winners and losers may act on different dimensions of aggressive behavior rather than simply producing opposite changes along a single axis. Importantly, a similar assumption of symmetry is evident at the molecular level, where winner and loser effects are often interpreted as opposite outputs of shared neuroendocrine and neurotransmitter systems [13]. Although these candidate-molecule approaches have provided important insights, they offer only a partial view of the molecular consequences of social experience. Consequently, transcriptomic approaches provide a broader perspective [14,15], but whether winner and loser effects are associated with distinct genome-wide molecular states remains unclear.

Transcriptomic studies of behavior have traditionally focused on differentially expressed genes (DEGs). Although this approach has advanced our understanding of the molecular underpinnings of social behavior [16,17], increasing evidence suggests that biologically meaningful differences may also reside in patterns of gene coexpression and network organization [18–22]. In sociogenomics [23–25], coexpression network analyses have elucidated behavior-dependent gene regulatory networks across a spectrum of social systems, from honeybees to vertebrates [20,26–30]. Such approaches may be particularly useful for studying winner–loser effects, as persistent behavioral states could be maintained through changes in transcriptomic organization even when large differences in individual gene expression are absent.

In this study, we combined behavioral analyses with whole-brain transcriptomic profiling and gene coexpression network analysis to investigate whether social competition produces distinct or shared persistent behavioral and molecular states in winners and losers of *Betta splendens*. To capture both the immediate and consolidated phases of the winner–loser effect, we collected whole-brain samples immediately after contests (A0; W0/L0) and five days later (A5d; W5d/L5d), alongside nonfighting controls (NF). Using this framework, we addressed three questions: (1) Do winner and loser experiences influence the same or different dimensions of aggressive behavior? (2) How do transcriptomic responses change from the immediate aftermath of social competition to the consolidated winner–loser state? and (3) What molecular features distinguish persistent winner and loser states?

*B. splendens* provides an ideal model for addressing these questions. Males of this fish engage in prolonged and highly structured contests that include a rich repertoire of quantifiable aggressive behaviors [31–33]. The winner effect typically persists for approximately five days, whereas the loser effect may last up to two weeks [34], providing a useful temporal window for examining both immediate and consolidated postfight states. The availability of its reference genome [35–37] further enables genome-wide investigations of the molecular consequences of social experience [38–40].

Our analyses revealed that winners and losers exhibited distinct behavioral outcomes and divergent patterns of transcriptomic network remodeling that emerged from a shared acute response. Together, these findings provide a system-level perspective on how social experience is translated into lasting behavioral and neurogenomic states.

## RESULTS

### Winner and loser effects diverge across distinct dimensions of aggressive behavior

To verify the persistence of winner and loser effects, each fish was tested with mirror assays before and after a size-matched contest followed by a 5-day rest period (Figure 1A; Methods). Aggressive behavior was compared among winners (W5d), losers (L5d), and NF controls using two composite measures: attack frequency (*AggFreq*), which captures aggressive output, and aggressive latency (*AggLat*), which captures the motivation to engage. Group differences were assessed with linear mixed-effects models (LMMs; *N* = 47; Table S1; Methods).

**Figure 1.**
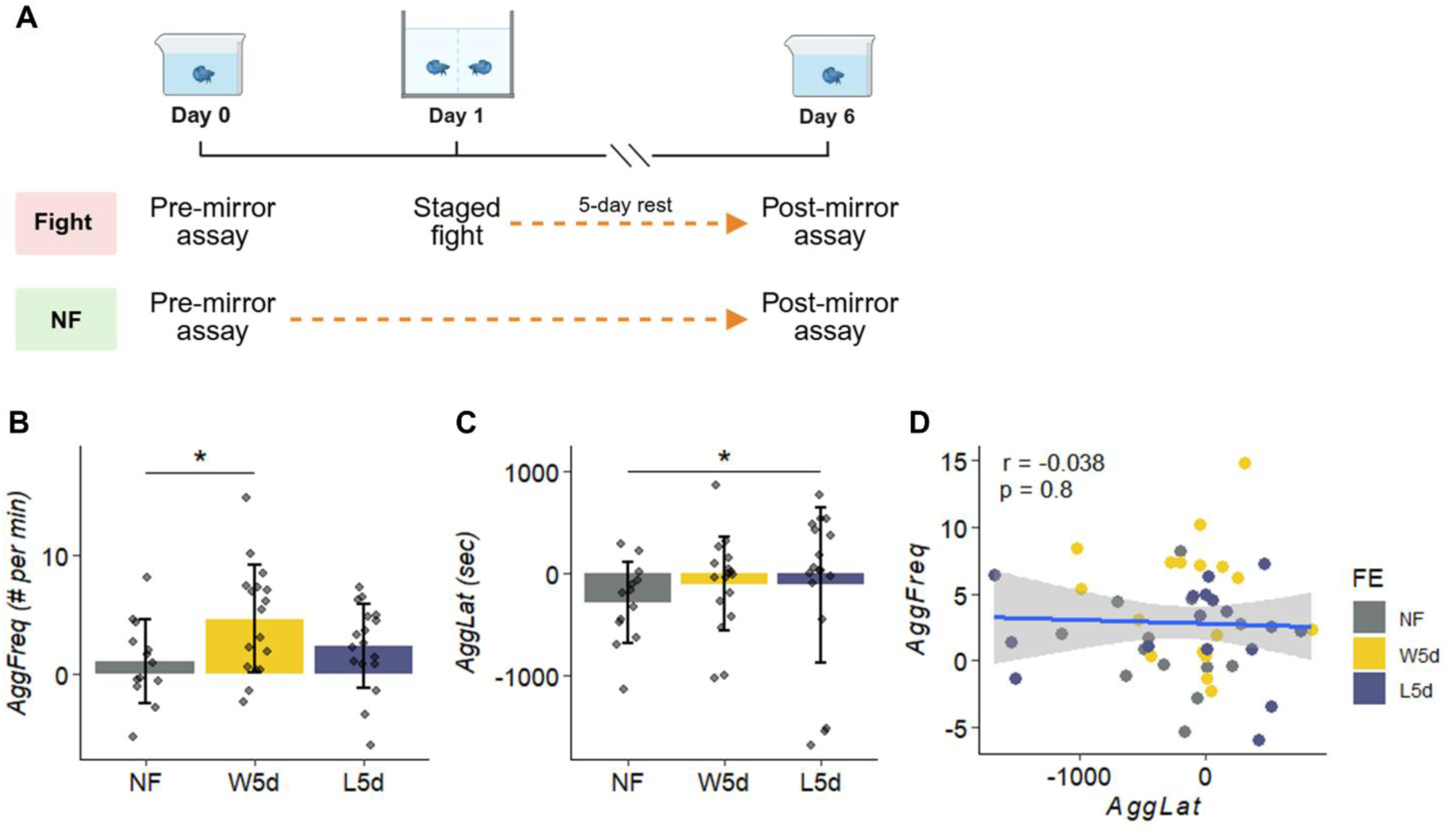
Winner and loser effects diverge across distinct dimensions of aggressive behavior. (A) Experimental design of the behavioral assays. Male *B. splendens* underwent a prefight mirror assay (Day 0), followed by either a staged fight or a nonfighting control (NF) treatment (Day 1) and a postfight mirror assay after a 5-day rest period (Day 6). Winners (W5d) and losers (L5d) were assayed at this later postfight time point. Created with BioRender.com. (B and C) Changes (post- minus preassay) in attack frequency (*AggFreq*, # per min) (B) and aggressive motivation (*AggLat*, sec; shorter latency indicates higher motivation) (C) across the NF, W5d, and L5d groups. *AggFreq* increased in W5d relative to that in NF, whereas *AggLat* differed in L5d relative to that in NF, indicating that the winner and loser effects manifested along separate behavioral dimensions. The bars show the mean ± SDs; the jittered points represent individual data points. The asterisks indicate significant differences from the NF controls, as determined by the linear mixed-effects models presented in Table 1 (B) and Table 2 (C), which accounted for preassay aggressive motivation/output and isolation duration, with pair identity included as a random intercept. (D) Correlations between changes in *AggLat* and changes in *AggFreq* across all individuals (Pearson correlation, with a fitted regression line and 95% confidence interval). \**p* < 0.05. *n* = 13 (NF), 17 (W5d), 17 (L5d); *n* = 47 total in (D). See also Table S1.

**Table 1.** Linear mixed-effects model predicting changes in attack frequency (*AggFreq*) in the mirror assays at the winner- (W5d) and loser-effect (L5d) states relative to nonfighting controls

| Fixed effects | Est. | S.E. | df | t | p |
| --- | --- | --- | --- | --- | --- |
| (Intercept) | 1.545 | 2.645 | 41.000 | 0.584 | 0.562 |
| Fighting experience |  |  |  |  |  |
| Winner | 3.532 | 1.591 | 41.000 | 2.221 | **0.032** |
| Loser | 1.261 | 1.594 | 41.000 | 0.791 | 0.433 |
| Level of gill cover erection count in preassay | -0.166 | 0.630 | 41.000 | -0.264 | 0.793 |
| Level of physical contact count in preassay | -0.594 | 0.659 | 41.000 | -0.901 | 0.373 |
| Isolation before experiment | 0.061 | 0.113 | 41.000 | 0.540 | 0.592 |

*The model was fitted by REML with Satterthwaite's method for degrees of freedom. Pair identity was included as a random intercept; NF controls were treated as independent observations. N = 47. A significant p-value is marked in bold.*

**Table 2.** Linear mixed-effects model predicting changes in aggressive initiation latency (*AggLat*) in the mirror assays at the winner- (W5d) and loser-effect (L5d) states relative to nonfighting controls

| Fixed effects | Est. | S.E. | df | t | p |
| --- | --- | --- | --- | --- | --- |
| (Intercept) | 185.508 | 275.096 | 38.545 | 0.674 | 0.5041 |
| Fighting experience |  |  |  |  |  |
| Winner | 161.764 | 169.614 | 41.746 | 0.954 | 0.3457 |
| Loser | 422.493 | 171.924 | 41.696 | 2.457 | **0.0182** |
| Level of latency time to the 1<sup>st</sup> aggressive acts in preassay | -507.844 | 87.632 | 41.924 | -5.795 | **<0.0001** |
| Isolation before experiment | 4.188 | 12.485 | 32.274 | 0.335 | 0.7395 |

*See Table 1 for model notes.*

Compared with the NF controls, winners showed a significant increase in *AggFreq* (Figure 1B; Table 1; *p* = 0.032), with a large effect size (*d* = 0.889, 95% CI [0.052, 1.726]), whereas no significant change was observed in *AggLat* (*p* = 0.346, *d* = 0.375; Table 2). Losers showed the opposite pattern, with a significant increase in *AggLat* (Figure 1C; Table 2; *p* = 0.018) and a large effect size (*d* = 0.979, 95% CI [0.154, 1.800]), but there was no significant change in *AggFreq* (*p* = 0.433, *d* = 0.317; Table 1). *AggFreq* and *AggLat* were not correlated (Figure 1D), supporting the interpretation that aggressive output and aggressive motivation represent distinct behavioral components. Taken together, the result of the behavioral analyses revealed dissociable effects of social experience: winning experience increased the frequency of aggressive actions, whereas losing experience prolonged attack latency. These findings suggest that, at the consolidated stage, the winner and loser effects selectively influence distinct components of aggressive behavior rather than representing opposite ends of a single aggression axis.

### Distinct postfight transcriptomic states emerge with structural winner–loser differences

Building on the observed behavioral dissociation between winners and losers, we investigated whether the winner–loser effect is associated with distinct brain transcriptomic states. RNA sequencing was performed on 28 whole-brain samples across different groups spanning two postfight stages (A0 and A5d) and NF controls, yielding 22,296 genes after filtering and normalization (Figure 2A; Methods).

**Figure 2.**
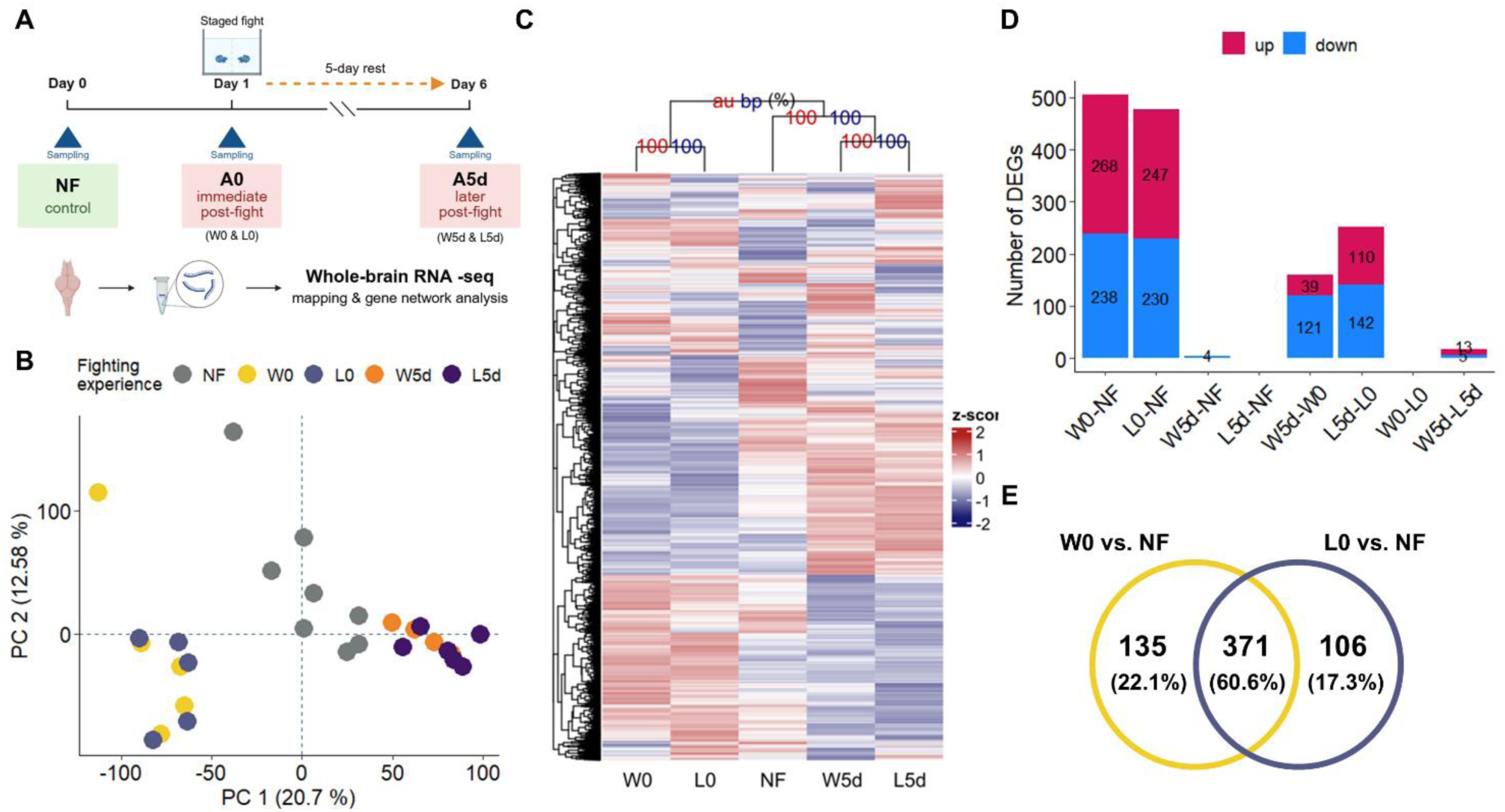
Distinct postfight transcriptomic states emerge with structural winner–loser differences. (A) Experimental design for whole-brain transcriptomic sampling. Whole-brain tissue was collected from nonfighting controls (NF), immediately after fighting (A0; winners, W0, and losers, L0), and after a 5-day rest period (A5d; winners, W5d, and losers, L5d), followed by RNA sequencing and transcriptional analysis, including gene network analysis. Created with BioRender.com. (B) Principal component analysis (PCA) of individual whole-brain transcriptomes across all groups. NF samples occupied an intermediate position between A0 and A5d samples along PC1, as confirmed by bootstrap-estimated centroid distances (see also Figure S2). (C) Hierarchical clustering and heatmap of group-averaged expression (*z* score) for 22,296 filtered genes across the NF, W0, L0, W5d, and L5d groups. The numbers at each node indicate approximately unbiased (AU, red) and bootstrap probability (BP, blue) values (%) from multiscale bootstrap resampling (2,000 replications). A0 samples clustered separately from NF and A5d samples, whereas winners and losers formed distinct subclusters within both A0 and A5d. (D) Number of differentially expressed genes (DEGs; FDR < 0.05) that were upregulated (magenta) or downregulated (blue) for each pairwise comparison. Comparisons with no bars indicate that no DEGs were detected in that direction. See also Table S2. (E) Venn diagram showing overlap of DEGs (up- and downregulated combined) between W0 vs. NF and L0 vs. NF comparisons, with percentages of the total union of DEGs indicated for each region. The majority of DEGs were shared between winners and losers in the immediate postfight state. The sample sizes for each group are listed in Table S1.

Principal component analysis (PCA), combined with bootstrap resampling of centroid distances and PERMANOVA, was used to characterize the overall transcriptomic variation across groups. The most prominent separation in PCA occurred between A0 (W0, L0) and the other groups along PC1 (20.7%; Figure 2B). Statistical analysis confirmed significant differences among all groups (PERMANOVA: *F_4,23_* = 3.627, *R^2^*= 0.387, *p* < 0.0001), while homogeneity of multivariate dispersions was consistent across groups (PERMDISP: *F_4,23_* = 0.680, *p* = 0.622). Within each postfight stage, winners and losers showed no clear separation in PCA space, with bootstrap-estimated centroid distances and pairwise PERMANOVA indicating substantial overlap at both A0 (*F_1,8_* = 0.267, *R^2^* = 0.032, *p* = 0.967) and A5d (*F_1,8_* = 0.781, *R^2^* = 0.089, *p* = 0.625; Figure S2). Notably, NF samples occupied an intermediate position between A0 and A5d along PC1, suggesting that a distinct longer-term molecular state emerged following the acute transcriptional response to fighting, rather than a simple reversion to baseline.

Hierarchical clustering provided a more nuanced perspective. While A0 samples were again distinctly separated from A5d and NF samples, winners and losers at both A0 and A5d formed separate clusters within each postfight stage, with full bootstrap support at both time points (Figure 2C). NF samples clustered with A5d rather than A0, indicating the appearance of a postfight molecular state that differs from the acute response. The separation of winners and losers within each stage, despite their proximity in the PCA space, suggests that their transcriptomic differences are structural rather than distance-based, reflecting a reorganization of gene expression relationships rather than substantial shifts in overall expression levels.

To further characterize these transcriptomic differences, we identified DEGs between groups using differential expression analysis (Figure 2D; Table S2). Immediately after the fight, both winners and losers exhibited transcriptional changes relative to the NF controls (W0 vs. NF: 506 DEGs; L0 vs. NF: 477 DEGs), with 371 DEGs shared between them (hypergeometric test, fold enrichment = 34.271, *p* < 0.0001; Figure 2E); however, no DEGs were detected between W0 and L0. By the later postfight stage, this signal had nearly resolved: only four DEGs were identified between W5d and NF, none between L5d and NF, and 18 between W5d and L5d. In contrast, substantial temporal remodeling was evident within each outcome group between A0 and A5d (winners: 160 DEGs; losers: 252 DEGs; Figure 2D). Together, these results suggest that the postfight stage, rather than contest outcome, is the primary driver of global transcriptomic variation.

### Shared acute neuroendocrine and signaling responses precede the winner- and loser-specific network remodeling

The discrepancy between the sparse set of DEGs at A5d and the persistent transcriptomic structure in the clustering suggests that differences between winners and losers may be encoded in the organization of gene coexpression rather than in individual expression levels. We therefore applied modular differential connectivity (MDC) analysis to characterize changes in the structure of gene coexpression networks across groups [22] (Methods). Using weighted gene coexpression network analysis (WGCNA), 22,296 genes from 28 samples were first partitioned into 43 modules (Table S3), and all the modules were subsequently subjected to MDC analysis (Table S4). Modules with significant batch effects on eigengene expression (FDR < 0.05; Methods) were subsequently excluded, and 20 modules were retained for interpretation (Figures 3, S3, and S4).

**Figure 3.**
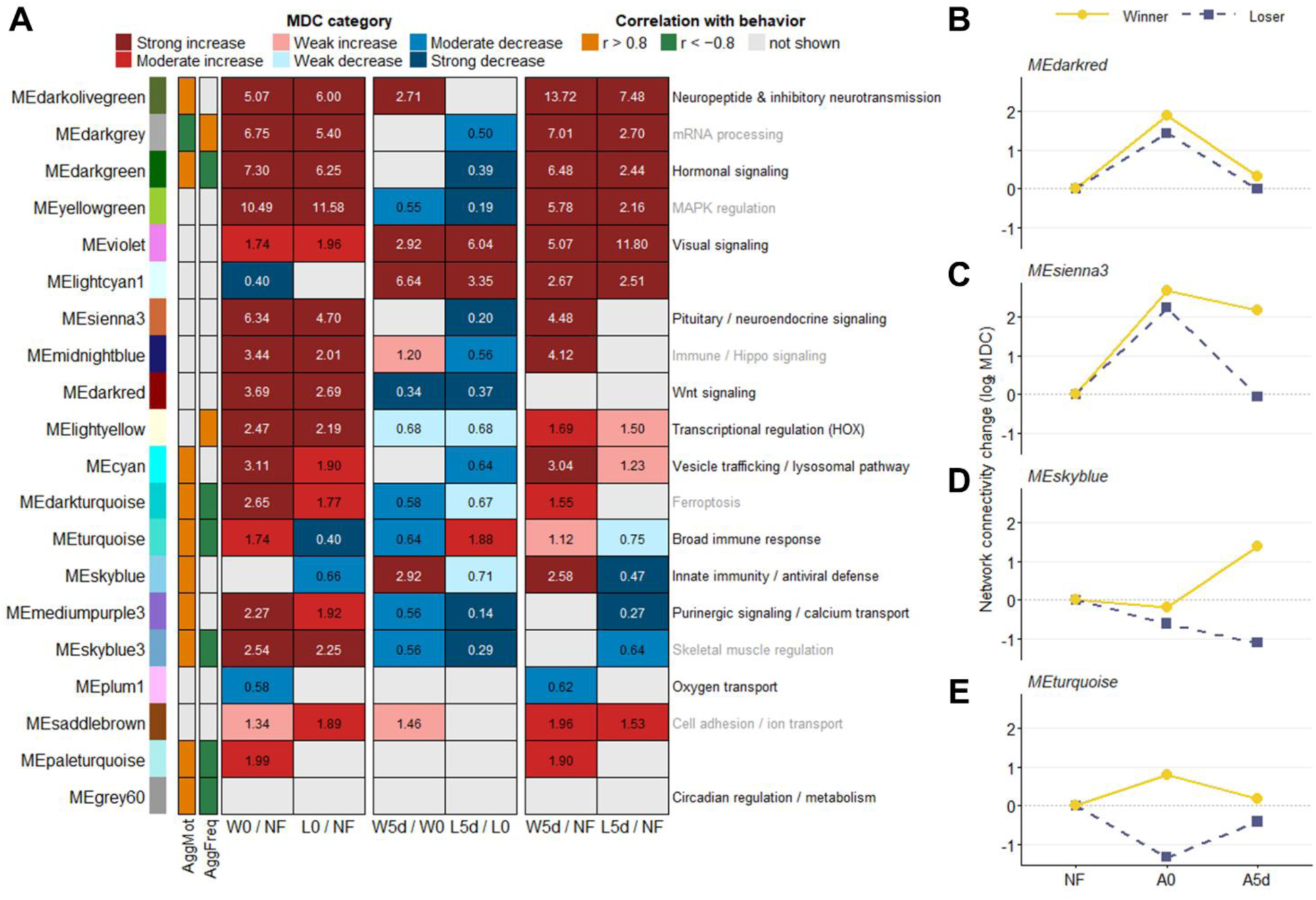
Winner and loser transcriptomic networks undergo asymmetric connectivity remodeling. (A) Modular differential connectivity (MDC) results for 20 coexpression modules retained after excluding modules with significant batch effects on eigengene expression (FDR < 0.05; Methods); the results for all 43 modules are shown in Figure S3. Heatmap cells indicate graded MDC classification (strong, moderate, or weak increase or decrease in connectivity) among comparisons with FDR < 0.05 (Methods). The values in the cells indicate the MDC values (see also Table S4). Left annotations indicate module eigengene correlations with aggressive motivation (*AggMot*, *-AggLat*) and aggressive output (*AggFreq*) (n = 3 group-level points; exploratory); orange, *r* > 0.8; green, *r* < -0.8; gray, |*r*| ≤ 0.8 (see also Figure S5 for complete *r* values). The right annotations list enriched functional terms (GO/KEGG; Table S5) in black (FDR < 0.05) or gray (raw *p* < 0.05). Module colors follow WGCNA naming conventions. Significance was assessed by a permutation test (Methods). (B–E) Network connectivity change (log_2_MDC, relative to NF) across the NF, immediate postfight (A0), and later postfight (A5d) states for winners (yellow, solid) and losers (purple, dashed) in four representative modules: MEdarkred (B), MEsienna3 (C), MEskyblue (D), and MEturquoise (E). The trajectories for all 20 modules are shown in Figure S4. The sample sizes for each group are listed in Table S1. See also Table S3.

At the acute stage, W0 and L0 exhibited highly similar patterns of network remodeling, sharing 14 significantly MDC-altered modules enriched for neuroendocrine (MEsienna3, MEdarkolivegreen), hormonal (MEdarkgreen), Wnt signaling (MEdarkred), and gene regulatory pathways (MEdarkgrey, MElightyellow), among others (Figures 3A, left, 3B, 3C, and S4; Table S5), with the notable exception of an immunity-associated module (MEturquoise), which already diverged in the direction between W0 and L0. Thus, despite their opposing contest outcomes, W0 and L0 exhibited a largely shared acute molecular response immediately after the fight.

As this shared acute response transitioned toward the consolidated winner–loser effect state (A0 to A5d), the gene coexpression network organization continued to remodel even though few DEGs remained detectable (Figure 2D). This transition involved both coordinated and divergent changes. Several modules shifted in the same direction in both winners and losers: connectivity increased in a visual signaling module (MEviolet) and in MElightcyan1 (including genes associated with transcriptional regulation, cellular metabolism, and mitochondrial homeostasis), whereas Wnt signaling (MEdarkred) and a purinergic signaling module (MEmediumpurple3) decreased. Moreover, a subset of modules began to diverge between winners and losers, such as neuroendocrine (MEsienna3) and hippo signaling (MEmidnightblue), indicating the emergence of winner- and loser-specific network reorganization (Figures 3A, middle, 3B, 3C, and S4; Table S5). These trajectories are consistent with the PCA patterns in which the NF samples occupied an intermediate position in the PCA space but were clustered with A5d rather than A0.

Together, these findings indicate a shared molecular starting point following social competition, with winner- and loser-specific transcriptomic states developing during the subsequent transition, suggesting that the postfight transcriptome evolves into a distinct long-term organizational state rather than simply returning to baseline.

### Asymmetric network reorganization underlies the established winner–loser effect

At the later postfight stage (A5d), winner- (W5d) and loser-specific (L5d) differences emerged and expanded against the shared network background established in the acute state. W5d showed increased coordination in modules associated with neuroendocrine signaling (MEsienna3) and innate immunity (MEskyblue), alongside reduced coordination in oxygen transport (MEplum1), whereas L5d exhibited reduced coordination in innate immunity (MEskyblue) and purinergic and calcium signaling (MEmediunpruple3), among others (Figures 3A, right, 3C, 3D, and S4; Table S5). The immunity-associated modules (MEskyblue and MEturquoise) showed opposing trajectories, increasing in winners but declining below baseline in losers, representing the most direct molecular contrast between the two effect states (Figures 3D and 3E). In particular, the divergence between the two groups regarding the immunity-associated module (MEturquoise) began immediately postfight. This pattern differs from the neuroendocrine (MEsienna3) and purinergic (MEmediumpurple3) modules, in which winners and losers did not diverge through opposite-direction changes. Instead, L5d became less coordinated than W5d related to NF, even though both groups initially experienced the same acute upregulation. Notably, these divergent changes occurred against the backdrop of persistent shared reorganization, with several acutely shared modules remaining elevated relative to baseline in both groups at the later postfight stage. Thus, winner–loser divergence did not reflect a symmetric, single-mechanism process of opposing changes but rather a structurally asymmetric reorganization of the network state (Figures 3A and S4; Table S5).

To determine whether these MDC alterations could be explained by the differential expression of module genes, we assessed DEG enrichment within each module using hypergeometric tests (Methods). Although several modules showed significant DEG enrichment at the acute and transition stages, DEGs consistently accounted for only a small fraction (<22.22%) of the total module membership (Figure S5A). No significant enrichment was detected at the later postfight state, despite the presence of multiple MDC-altered modules at this stage (Figure S5B). Notably, standard DEGs alone would have concluded that winners and losers were largely indistinguishable at the later postfight stage, whereas MDC analysis revealed persistent, structured differences in network coordination. This contrast highlights that the molecular signature of the established winner–loser effect is captured by coexpression network reorganization rather than by individual gene expression.

### Behavioral variation corresponds with transcriptomic network organization

Having established that winner–loser differences reside primarily in network organization, we next asked whether these networks were also associated with the behavioral variations observed in the winner–loser effects. At minimum, the correlation of module eigengenes with *AggMot* (-*AggLat*) and *AggFreq* across groups revealed 12 modules with strong correlations (|*r*| > 0.8) with at least one behavioral measure (Figure S6; as annotated in Figure 3A; Methods). Notably, these modules were dissociated along the two behavioral dimensions, and this dissociation aligned with whether the modules overlapped with the previously identified winner- or loser-specific MDC modules.

Six modules overlapping with the winner- or loser-specific MDC modules at the later postfight stage were primarily positively associated with *AggMot*, and four of these modules were also negatively correlated with *AggFreq* (as annotated in Figure 3A). Among these, the immune-related modules showing the most pronounced divergence between winners and losers according to MDC analyses (MEturquoise, MEskyblue) were positively correlated with *AggMot*, indicating that immune network coordination was most consistently correlated with the suppression of the behavioral dimension in losers.

The remaining modules did not overlap with winner- or loser-specific MDC modules at the later postfight stage and were divided on the basis of behavioral dimensions: those positively associated with *AggMot* (MEgrey60, MEdarkgreen, MEcyan, and MEdarkolivegreen) were enriched for metabolic and neuromodulatory functions (Table S5). Conversely, modules positively associated with *AggFreq* (MEdarkgrey, MElightyellow) exhibited opposite correlation directions and were enriched for gene regulatory functions. It is important to note that, because the behavioral and transcriptomic data were obtained from separate individuals, these associations represent group-level patterns rather than individual-level correlations and should be treated as exploratory findings. Collectively, these patterns suggest that *AggMot* and *AggFreq* correspond to partially distinct transcriptomic networks, paralleling the behavioral dissociation observed between winner and loser effects.

## DISCUSSION

This study reveals the behavioral and transcriptomic asymmetries that underlie the winner–loser effect in *B. splendens*. Behaviorally, winning experience selectively increased attack frequency, whereas losing experience primarily elevated aggressive latency, indicating that the winner and loser effects influence distinct aspects of aggressive behavior rather than producing inverse changes along a singular aggression continuum. At the transcriptomic level, both winners and losers initially exhibited acute neuroendocrine and signaling responses, subsequently diverging in patterns of network remodeling specific to each. The loser-effect state was characterized by widespread immune network discoordination, whereas the winner-effect state demonstrated sustained coordination of neuroendocrine and immune defense networks. Notably, these persistent differences were more pronounced in the organization of gene coexpression networks than in enduring differential gene expression, implying that the winner–loser effect is predominantly encoded by the reorganization of transcriptomic networks following an initial shared response to social conflict.

### Behavioral asymmetry of the winner–loser effect

At the behavioral level, the winner and loser effects clearly differed: winning selectively increased attack frequency (*AggFreq*), whereas losing selectively prolonged aggressive latency (*AggLat*), with neither experience significantly affecting the other dimension. Previous studies have seldom distinguished between the motivational and output components of aggression; when both measures were recorded, the results were generally interpreted as reflecting a common aggressive strategy [1,41]. The present results suggest that this assumption may require re-evaluation.

A potential concern is that the 5-day intercontest interval may have allowed the loser effect on attack frequency to decay before assessment. Experience-dependent behavioral effects are known to diminish over time [42,43], and the loser effects on attack frequency have been documented at shorter intervals [1]. Indeed, previous research on mangrove killifish (*Kryptolebias marmoratus*) has shown that the apparent persistence of winner and loser effects varies with the behavioral measure employed, with different metrics yielding divergent conclusions regarding the duration of each effect [11]. However, both behavioral dimensions were subjected to the same 5-day interval, thereby experiencing identical decay pressures. The fact that *AggLat* remained significantly elevated among losers, whereas *AggFreq* did not, suggests that these two dimensions differ in their persistence rather than merely in their direction. The persistence of the winner effect on *AggFreq* after the same interval further underscores an asymmetry in the temporal dynamics between the two effects [1]. The near-zero correlation between *AggFreq* and *AggLat* (*r* = −0.038), together with their differing sensitivities to social experience, supports the interpretation that aggressive output and aggressive motivation represent distinct behavioral components shaped by partially independent mechanisms.

This behavioral dissociation also has evolutionary implications. The selective increase in attack frequency after winning may facilitate success in future encounters [44], whereas the persistent increase in aggressive latency in losers may serve as an adaptive strategy to reduce the risk of further injury during recovery [1]. Evidence that the duration of the loser effect is heritable and can evolve independently of baseline fighting ability [45] further supports the view that the winner and loser effects represent partially independent evolutionary outcomes rather than opposite ends of a single selective axis.

### Social competition elicits a shared acute neurogenomic response

Despite the pronounced behavioral differences observed between winners and losers, the persistent transcriptomic divergence at the level of individual gene expression remains limited. PCA revealed that the postfight stage, rather than the contest outcome, was the dominant source of transcriptomic variation. This pattern is consistent with the results of the DEGs analyses, in which winners and losers shared hundreds of DEGs immediately after contests but exhibited no detectable transcriptional differences between outcomes. A comparable dissociation between behavioral divergence and differential gene expression has recently been reported in zebrafish, where winners and losers exhibited marked behavioral differences but limited transcriptomic differentiation at the DEG level [46]. Broadly speaking, accumulating evidence indicates that meaningful biological differences between behavioral states may be embedded in network organization, even when few genes are differentially expressed [18,19,47].

Immediately after the fight, winners and losers exhibited a highly similar pattern of transcriptomic network remodeling: MDC analysis similarly revealed extensive remodeling of neuroendocrine, hormonal, Wnt signaling, and gene regulatory modules, despite the limited identification of DEGs noted above. This observation indicates that the initial molecular responses to social competition are largely independent of contest outcome. Such acute responses align with the conserved neuromolecular toolkit activated by acute social challenges across vertebrate species[20,48–50], and are consistent with findings in paper wasps, zebrafish, and *B. splendens*, where both winners and losers display comparable transcriptomic profiles immediately postinteraction, diverging over subsequent hours or days [16,38,51]. Collectively, these observations suggest that social competition first elicits a common neurogenomic state reflecting an acute response to social challenge rather than the eventual winner-or-loser phenotype, indicating an evolutionarily conserved molecular reaction to social conflict across distantly related species.

### Winner and loser states emerge through distinct patterns of network remodeling

Although winners and losers exhibited similar acute transcriptomic responses, they progressively diverged through asymmetric trajectories in terms of both temporal dynamics and functional composition. This difference was not primarily reflected at the level of differential gene expression. Instead, hierarchical clustering consistently separated winners from losers despite the limited number of persistent DEGs, suggesting that the difference in transcriptome between winners and losers was organizational rather than expression level [52]. Winners showed sustained increases in network coordination across modules associated with neuroendocrine function (MEsienna3) and intracellular trafficking (MEcyan), whereas losers showed persistent discoordination in the purinergic signaling module (MEmediumpurple3). Both the purinergic signaling pathway and the neuroendocrine signaling pathway are known regulators of social behavior, aggression, and physiological responses to challenge [50,53–55]. The stronger network coordination of intracellular trafficking in winners than in losers is consistent with a greater capacity for synaptic plasticity and neural adaptation following victory [2,56,57].

Among these modules, the purinergic signaling module exhibited a strong positive correlation with *AggMot* across groups, progressively diverging between winners and losers. It remained significantly below baseline solely in losers within the established states, suggesting that its persistent discoordination in the loser-effect state may be related to the reduced aggressive motivation observed in losers. Although this correlation is based on group-level means and should be treated as an exploratory finding, purinergic signaling represents a candidate pathway for future mechanistic investigation. The neuroendocrine module followed a divergent trajectory: winners maintained coordinated network activity while losers returned to baseline but showed no strong correlation with either behavioral measure. Previous studies have largely emphasized hormones and neurotransmitters as candidate mediators of winner and loser effects [13,43,48,58]. The present results extend this view by showing that winner and loser effects involve coordinated changes in network connectivity that are not captured by individual candidate molecules or by the differential expression of single genes. For consolidated behavioral states in which few genes exhibit sustained differences in expression, coexpression network analysis may be necessary—not merely complementary—to reveal the underlying molecular organization. Together, these observations suggest that persistent winner and loser phenotypes emerge primarily through transcriptomic network reorganization rather than long-lasting changes in individual gene expression. Circuit-level studies have shown that social experience can remodel specific neural pathways, such as the thalamo–prefrontal circuit, which reinforces social dominance, and the habenular subregions, which govern conflict resolution [59,60]. The whole-brain network reorganization described here offers a complementary, brain-wide perspective on these processes, and linking network-level coordination to such defined circuits represents an important direction for future work.

### Immune dysregulation as a potential molecular signature of the loser effect

Acute stress following social conflict is typically expected to be accompanied by an upregulated immune response, reflecting an adaptive physiological reaction to the risk of injury and increased pathogen exposure during contests [61,62]. In this study, the neurotranscriptomic profile of the winner effect revealed a sustained increase in the coordination of immune-related networks following the fight, which was consistent with the expected increase in the immune response. The loser-effect state, however, did not simply fail to exhibit the expected response; instead, immune network coordination decreased significantly below the nonfight baseline. The most prominent feature of the loser-effect state identified in this study was the discoordination of immune-related gene networks, a pattern that first emerged in the acute stage and persisted into the consolidated state. The module associated with a broad immune response (MEturquoise) showed divergence in network coordination between winners and losers from the acute stage. In contrast, the module linked to innate immunity and antiviral defense (MEskyblue) exhibited a more progressive divergence that intensified throughout the transition period, ultimately leading to markedly opposing coordination patterns between winners and losers in the final stage. This pattern of immune network discoordination in losers is unlikely to be attributable to physical injuries from a fight. Even if losers sustain greater physical damage during contests, immune activation driven by peripheral injury would presumably increase immune network coordination in both groups. Moreover, research in fish has indicated that peripheral immune activation does not consistently or immediately produce changes in gene expression within the brain, with responses varying across distinct brain regions and emerging at different time points [63]. Consequently, the peripheral wound response alone is unlikely to account for the opposing trajectories of immune networks observed immediately after the fight.

Winners and losers showed opposing coordination of immune networks, paralleling the divergent immune profiles reported between dominant and defeated individuals in mammals [47,64–67]. This correspondence suggests that immune responses to social competition may be evolutionarily conserved. From an evolutionary perspective, this bidirectional immune regulation may reflect an adaptive strategy shaped by the contrasting demands of social competition [68]. For winners, social dominance confers access to resources, territory, and mating opportunities, but also entails increased social contact and pathogen exposure; maintaining strong immune capacity may therefore be advantageous [66,67]. Losers, which have been reported to experience social withdrawal following defeat [1,69], may correspondingly reduce pathogen exposure and reallocate immune resources toward wound healing [67,70,71]. This behavioral and immunological pattern is consistent with the social competition hypothesis of depression [72], in which withdrawal from social interaction represents an adaptive response to defeat. Notably, immune dysregulation in mammals has typically been reported following repeated social defeat [67], whereas the network-level discoordination observed here emerged after a single contest, suggesting that a single defeat experience may be sufficient for persistent immune network reorganization to emerge. Although our behavioral assays do not directly assess depressive states [73], the molecular resemblance to mammalian immune dysregulation suggests a shared neuroimmune response rather than clinical equivalence. While this bidirectional relationship has been documented in mammals at the level of immune function and cytokine profiles, the opposing coordination of the antiviral defense network in winners and losers reported here provides, to our knowledge, the first evidence of this pattern at the level of gene coexpression networks in a nonmammalian vertebrate.

## Conclusion

In summary, under near-natural ecological conditions, this study reveals that the winner–loser effect in *B. splendens* operates through behavioral and neurogenomic asymmetries rather than opposing changes along a single axis. Winning selectively increased aggressive output, whereas losing selectively reduced aggressive motivation, indicating that the two effects influence distinct behavioral dimensions. At the transcriptomic level, social competition initially elicited a shared acute neuroendocrine response and then diverged into winner- and loser-specific patterns of network remodeling across functionally distinct networks, including those associated with immune, neuroendocrine, and purinergic signaling. These networks diverged in different ways and on different timescales, with immune network discoordination emerging as the primary molecular signature of the loser-effect state. These findings provide network-level neurotranscriptomic evidence from a nonmammalian vertebrate that is consistent with the proposed evolutionarily conserved relationship between social defeat and immune dysregulation, and with depression-related immune signatures reported in mammals.

### Limitations of the study

While the present study provides new insights into the neurogenomic basis of the winner–loser effect, several limitations warrant consideration. First, the strict inclusion criteria used to obtain unambiguous winner and loser outcomes yielded a relatively small sample size, particularly for the W5d group (N = 4; Methods). Although key modules remained stable across sensitivity analyses (Methods), findings specific to this group should be interpreted with caution. Second, coexpression network analysis was performed on normalized expression values without prior batch correction, as applying batch correction before WGCNA risks removing biologically meaningful coexpression structures. Modules significantly associated with sequencing batches were subsequently excluded (see Methods), which may also have reduced the recovery of biological signals. Finally, whole-brain transcriptomics lacks the spatial and cellular resolution needed to identify the specific neural circuits and cell populations underlying the observed network changes. Future studies incorporating additional postfight time points, spatially resolved or single-cell transcriptomic approaches, and validated affective state paradigms in *B. splendens* would help resolve the full temporal dynamics of network remodeling and test the mechanistic roles of the candidate pathways identified here.

## MATERIALS AND METHODS

### Experimental model and study participant details

Male adult *Betta splendens* [31] were sourced from a commercial supplier and imported from Thailand. Upon arrival, each fish was housed individually in 600-mL glass bottles and quarantined for three days before experimental procedures. Opaque dividers prevented visual contact between the fish. They were maintained at 27 ± 2°C under a 12:12-h light–dark cycle and fed a commercial diet (Hikari Tropical Vibra Bites, Kyorin Food Industries, Ltd., Japan) twice daily. After quarantine, the fish underwent an additional isolation period before the experiment (total isolation time: 8.4 ± 4.36 days; range: 3–18 days), with this duration serving as a covariate in the behavioral analyses [1]. The average length and weight of the fish were 4.9 ± 1.17 cm and 2.04 ± 0.49 g, respectively. The fish were paired by color morph and body size. Each pair was then randomly assigned to either the behavioral experiment or the brain transcriptomics experiment; nonfighting (NF) controls were also randomly assigned to one of the two experiments. All procedures were approved by the Institutional Animal Care and Use Committee (IACUC) of National Cheng Kung University, Taiwan (approval no. 110116).

### Experimental design

This study involved two experimental components: a behavioral component and a brain transcriptomics component. Fish were assigned to either a fight paradigm or an NF control condition. Fight-experienced fish underwent a sequential screening procedure; full sample attrition details are provided in Figure S1. Throughout the manuscript, samples are grouped by postfight stage: nonfighting controls (NF), immediately postfight (A0), and 5 days postfight (A5d). Within each postfight stage, winners and losers are denoted W0/L0 (A0) and W5d/L5d (A5d), respectively.

Fighting trials were conducted in transparent glass tanks (20 × 12 × 15 cm; 3 L of water), divided by an opaque partition. Each fish was placed in a separate compartment and allowed to acclimate for 15 minutes without visual contact before the partition was removed to initiate the trial. All behavioral assessments were video-recorded using a digital camcorder (JVC GZ-F170). Fighting trials were conducted between 10:30 and 16:30 (UTC+8).

Male *B. splendens* fights involve a repertoire of aggressive behaviors ranging from noncontact displays (e.g., gill cover erection, lateral displays) to escalating physical contact (tail beats, bites, and mouth locking, in order of increasing intensity), culminating in a clear winner–loser outcome characterized by persistent chasing and fleeing [32,33]. Not all fights escalate to mouth locking; therefore, only trials in which mouth locking was observed were retained to ensure that unambiguous, fully escalated fights were included in the analysis. Trials in which a winner–loser outcome was established within 30 minutes were excluded because such rapidly resolved fights may reflect large intrinsic asymmetries in fighting ability rather than a genuine contest and may be insufficient to reliably induce winner and loser effects [1]. Trials exceeding two hours without a clear outcome were also excluded.

For the behavioral component, each fish underwent a mirror assay on Day 0 (prefight baseline), participated in a staged fight on Day 1, and underwent a second mirror assay on Day 6 (postfight assessment) after a 5-day rest period in individual housing without visual contact with conspecifics. NF controls were maintained under identical housing conditions and subjected to the same mirror- assay schedule, without prior fighting experience (Figure 1A).

With respect to the brain transcriptomics component, the fish participated in a staged fight on Day 1 using the same protocol. W0 and L0 samples were collected within three minutes of the fight’s conclusion, between 10:30 and 12:30 (UTC+8). W5d, L5d, and NF samples were collected on Day 6 between 12:30 and 13:30 (UTC+8). Sacrifice time points were deliberately aligned across groups to control for circadian variation in gene expression (Figure 2A).

After applying these exclusion criteria, the final behavioral sample comprised 47 individuals (W5d = 17, L5d = 17, NF = 13; Table S1). For the brain transcriptomics component, further sample exclusions due to tissue collection and RNA quality are described in the Brain sampling and RNA extraction section.

### Behavioral assessment: Mirror assay

Aggressive behavior was assessed using a mirror assay following Balzarini et al. [74]. Each fish was placed individually in a test tank (13 × 35 × 17 cm, 3 L) with an acrylic mirror (11 × 16 cm) mounted on one of the side walls. An opaque partition covered the mirror during a 5-minute acclimation period and then was removed to initiate the 20-minute trial. All trials were video-recorded (JVC GZ-F170) and coded post hoc in BORIS v.9.7.13 [75] by a naïve observer who was blinded to group identity and fighting outcome.

Gill cover erection and physical contact (tail beats and bites combined) were each scored as total event counts and latency to first occurrence. Since threatening displays and physical attacks may reflect distinct mechanisms along the aggression escalation cascade [1,76], changes in attack frequency (*AggFreq*, expressed as counts per minute over the 20-minute trial) and aggressive latency (*AggLat*) between post- and prefight assays were summed across both behavior types to form composite measures of aggressive output and motivation, respectively. The two composite measures were largely independent of each other (*r* = −0.038, *p* = 0.800), supporting their treatment as distinct behavioral dimensions. Although the two components of each composite differed in variance, sensitivity analyses using *z* score-standardized composites yielded virtually identical results (*AggFreq*: winner effect *p* = 0.037, *d* = 0.866; *AggLat*: loser effect *p* = 0.020, *d* = 1.017), confirming robustness to the choice of standardization. Statistical analyses are described in the Quantification and statistical analysis section.

### Brain sampling and RNA extraction

Fish were anesthetized by immersion in MS-222 (168 mg/L; APExBIO, CAS no. 886-86-2) for approximately 3 minutes and immediately submerged in liquid nitrogen to minimize RNA degradation [77]. The samples were stored at −80°C until dissection. For dissection, frozen samples were placed in ice-cold 1X phosphate-buffered saline (PBS) until the tissues had thawed and softened. The whole brain was rapidly dissected and immediately transferred to a 1.5 mL RNase-free tube containing RNAlater reagent (Invitrogen and QIAGEN) and stored at 4°C for up to one week before RNA extraction.

Total RNA was extracted using the Invitrogen PureLink RNA Mini Kit (cat. no. 12183018A) with on-column DNase treatment (PureLink DNase). Briefly, each brain was homogenized in 400 µL of lysis buffer containing 1% 2-mercaptoethanol and 600 µL of TRIzol LS Reagent (Invitrogen, cat. no. 15596018) using a BioMasher disposable homogenizer (Nippi, Tokyo, Japan). Following homogenization, 100 µL of 1-bromo-3-chloropropane (BCP) was added, and the mixture was vortexed and centrifuged at 12,000 × g for 10 minutes at 4°C. An equal volume of 70% ethanol was added to the aqueous phase, and RNA was purified using spin columns per the manufacturer’s instructions. RNA quality and concentration were assessed using an Agilent 4200 TapeStation with High Sensitivity RNA ScreenTape (Agilent Technologies); samples with an RNA integrity number (RIN) < 6.5 were excluded. Six samples were lost during dissection, and four additional samples were excluded because of RNA quality failure, yielding a final transcriptomic sample of 28 individuals (W0 = 5, L0 = 5, W5d = 4, L5d = 6, NF = 8; Figure S1; Table S1).

### RNA sequencing and bioinformatic processing

Library preparation and Illumina NovaSeq sequencing (2×150 bp paired-end, strand-specific with polyA selection, ∼35 million reads per sample) were conducted by Azenta Life Sciences (Tokyo, Japan) in multiple batches; the sequencing batch was treated as a covariate in downstream analyses as described in the relevant sections below.

Raw reads were preprocessed using the following pipeline: adapter sequences were trimmed with Cutadapt v.3.5 [78]; poly(A) tails were removed using fastx_clipper; low-quality bases were trimmed using fastq_quality_trimmer (parameters: -t 20 -l 30 -Q 33); reads failing quality thresholds were discarded using fastq_quality_filter (parameters: -q 20 -p 80 -Q 33), all from FASTX-Toolkit v.0.0.14; unpaired reads were removed with Trimmomatic v.0.39 [79]; and rRNA, globin, and phiX sequences were removed with Bowtie2 v.2.4.4 [80], followed by the recovery of unmapped reads with bam2fastq v.1.1.0. For each sample, 30 million read pairs were subsampled prior to genome mapping. Reads were mapped to the *B. splendens* reference genome (NCBI RefSeq GCF_900634795.4) using HISAT2 v.2.2.1 [81] with the --dta flag, and uniquely mapped reads were extracted using SAMtools v.1.13 (parameters: view -q 4) [82]. Read counts per gene were quantified at the exon level using featureCounts v.2.0.3 (parameters: -t exon -g gene_id) [83].

Genes with low expression were filtered using the filterByExpr function in edgeR v.4.8.2 [84], retaining 22,296 genes. Filtered counts were normalized using the trimmed mean of M-values (TMM) method. Batch-effect correction strategies are described in the relevant sections below. All downstream analyses were performed in R v.4.5.1.

### Principal component analysis and hierarchical clustering

Batch-corrected, TMM-normalized expression values were obtained by applying removeBatchEffect (limma v.3.66.0) to log-CPM values, using the sequencing batch as the correction variable and a design matrix that included the fighting experience group (W0, L0, W5d, L5d, and NF) as a protection variable to prevent the removal of group-level expression differences. The resulting matrix served as input for all subsequent multivariate analyses.

Principal component analysis (PCA) was performed on the batch-corrected expression matrix using prcomp (R base package) with centering and scaling. Group-level differences in PCA space were assessed using PERMANOVA (adonis2 from vegan v.2.7.3, 9999 permutations) on Euclidean distances computed from the first 10 principal components, with homogeneity of group dispersions confirmed using betadisper (9999 permutations). Pairwise PERMANOVA was performed between winner and loser groups at each postfight stage. To estimate the stability of the group centroids, a bootstrap procedure (R = 1000; seed = 111) was applied: for each resample, PCA was recomputed, the PC directions were aligned with the observed solution by sign correction, and pairwise Euclidean distances between group centroids in the PC1–PC2 space were calculated. The bootstrap distributions of the centroid distances are summarized as medians and 95% confidence intervals (Figure S2).

For hierarchical clustering, group-mean expression profiles were computed across all 22,296 genes, and *z* scores were calculated per gene. Hierarchical clustering with bootstrap support was performed using pvclust v.2.2-0 [85] with correlation distance, average linkage, and 2000 bootstrap replicates (seed = 328). Clusters with approximately unbiased (AU) bootstrap support ≥ 95% were considered well supported. The resulting dendrogram was used to order columns in a heatmap generated with ComplexHeatmap v.2.26.1 [86]; rows were clustered using default settings (Euclidean distance, complete linkage).

### Coexpression network construction and module differential connectivity analysis

Coexpression network analysis was performed using weighted gene coexpression network analysis (WGCNA) v.1.74 [87] on TMM-normalized log-CPM values before batch correction, following the official WGCNA tutorial with default parameters. The soft-thresholding power was set to 8, the lowest power that achieved *R^2^* ≥ 0.80 for the scale-free topology model fit (*R^2^* = 0.882) while maintaining adequate mean connectivity (*mean.k* = 87.3). Network construction yielded 43 coexpression modules (Table S4).

MDC analysis was performed on all 43 modules, following Fang et al. [22], to quantify changes in intramodular coexpression connectivity between groups. For each pairwise comparison, MDC was calculated as the ratio of the sum of intramodular connectivity between the two groups. Statistical significance was assessed by a permutation test (10,000 permutations, seed = 1000) with FDR correction across modules. The significant results were further classified into six grades on the basis of the magnitude of MDC: strongly increased (MDC > 2), moderately increased (MDC 1.5–2), weakly increased (MDC 1–1.5), weakly decreased (MDC 1/1.5–1 [≈0.67–1]), moderately decreased (MDC 0.5–1/1.5 [≈0.5–0.67]), and strongly decreased (MDC < 0.5). Six pairwise comparisons were performed: W0/NF, L0/NF, W5d/W0, L5d/L0, W5d/NF, and L5d/NF (Figure S3; Table S4).

Given that MDC analysis estimates intramodular connectivity within each group independently and that the W5d group comprises only four samples, which may influence the stability of coexpression modules, WGCNA was repeated with soft-thresholding powers of 6, 9, and 12 to assess whether module definitions remained consistent across different parameter settings (Table S6). Ninety percent of the modules at a power of 9 had a Jaccard similarity ≥ 0.5 relative to those at a power of 8, whereas lower stability was observed at a power of 6 (67%) and a power of 12 (76%), which is consistent with the anticipated sensitivity of smaller modules to the choice of power. Modules central to the primary findings exhibited consistently high Jaccard similarity across powers of 6, 9, and 12, except for MEskyblue at a power of 6, which reflects its relatively small size and susceptibility to network resolution. Collectively, these findings support the selection of a power of 8 and suggest that the module definitions that underpin the principal conclusions are both stable and dependable.

To exclude modules confounded by sequencing batch effects, a linear model was fitted for each module eigengene (ME), with the fighting experience group and batch as predictors (ME ∼ group + batch). The batch term was evaluated by type I ANOVA with FDR correction across all 43 modules. Modules with an FDR < 0.05 for the batch term were excluded, yielding 20 modules retained for interpretation (Figure 2).

### Differential gene expression analysis

DEGs were identified using edgeR v.4.8.2 [84]. A design matrix was constructed to include the sequencing batch as a covariate and the fighting experience group (W0, L0, W5d, L5d, and NF) as the main effect. Dispersion was estimated using estimateDisp with robust estimation, and a quasilikelihood negative binomial generalized linear model was fitted using glmQLFit [88]. Pairwise comparisons were performed using glmQLFTest, and genes with a false discovery rate (FDR) < 0.05 were considered differentially expressed. The statistical significance of the overlap between W0 and L0 DEGs was assessed using a hypergeometric test implemented via the phyper function in R, with all expressed genes (*N* = 22,296) as the background set. The magnitude of over-representation was quantified by fold enrichment, calculated as the ratio of the observed overlap proportion to the expected proportion under random chance.

### Module–DEG Enrichment Analysis

To assess whether coexpression modules were enriched for DEGs from each pairwise comparison, a hypergeometric test was performed using the phyper function in R. DEGs (FDR < 0.05) from each comparison were tested for enrichment against gene member lists from the 20 batch- unconfounded coexpression modules, with the total number of detected genes (*N* = 22,296) serving as the background. For each module, the probability of observing an overlap equal to or greater than that detected by chance was calculated (upper-tail, one-tailed), and the resulting *p* values were corrected for multiple comparisons using the Benjamini–Hochberg procedure (FDR < 0.05).

### Module eigengene–behavior correlation

To examine the correspondence between neurogenomic states and behavioral phenotypes, Pearson correlations were computed between group-mean module eigengene values and group-mean behavioral metrics. Analyses were restricted to the NF, W5d, and L5d groups, as behavioral postfight assessments were available only for these groups. Group means were calculated for each module eigengene and for *AggFreq* and *AggMot* (−*AggLat*, where higher values reflect greater aggressive motivation) [89]. Pearson correlation coefficients were computed between each module eigengene and each behavioral metric across the three group means. Given that correlations were computed across only three group means, these analyses are intended to be exploratory and descriptive rather than inferential.

### Gene Functional Enrichment Analysis

Gene Ontology biological process (GO-BP) and Kyoto Encyclopedia of Genes and Genomes (KEGG) pathway enrichment analyses were performed using the Database for Annotation, Visualization and Integrated Discovery (DAVID) Bioinformatics Resources (DAVID Knowledgebase v2025_2; https://davidbioinformatics.nih.gov/) [90,91], with *B. splendens* as the background species. The results for which the FDR was < 0.05 were considered significant; when no terms met this threshold, the results with *p* < 0.05 were reported as suggestive.

### Quantification and statistical analysis

Behavioral data were analyzed using linear mixed-effects models (LMMs) fitted by restricted maximum likelihood (REML) with Satterthwaite’s method for degrees of freedom, implemented in lmerTest v.3.2-1 [92]. The fighting experience group (NF, W5d, and L5d) was the main fixed effect in all the models. Prefight behavioral covariates were included to control for individual baseline differences in aggressive tendency and metabolic capacity. Because the raw behavioral counts showed high interindividual variability, each covariate was binned into five ordinal levels prior to model fitting: the prefight gill cover erection count and physical contact count were coded in ascending order (higher levels reflecting greater aggressive output), and the prefight latency to first aggressive action and prefight surface breathing count were coded in descending order (higher levels reflecting greater aggressive motivation and greater aerobic capacity, respectively). Each model included the corresponding prefight behavioral covariate(s) and total isolation duration as a continuous covariate [1,93]. Pair identity was included as a random intercept to account for the nonindependence of paired individuals; NF controls were treated as independent observations, as they had no fighting partner. Effect sizes (Cohen’s *d*) and 95% confidence intervals were calculated using eff_size (emmeans v.2.0.3) [94], with the residual standard deviation of each model as the pooled standard deviation estimate.

## ACKNOWLEDGEMENT

This work was partially supported by the laboratory of N.O. (Kitasato University), funded by Tsumura & Co. (Japan), and by the laboratory of H.-V.W. (National Cheng Kung University). We thank Takefumi Nakazawa and H. Sunny Sun (National Cheng Kung University) for providing the experimental space, equipment, and other practical support during this study. M.-T.C. was supported by a Scholarship for Outstanding Doctoral Students from the Ministry of Science and Technology, Taiwan. We thank Hao-Bin Chen and Wei-Jen Lee for assisting with the sample collection, including conducting and video-recording behavioral trials. We thank I-Wen Chen, Chia-Hsuan Wei, and the Consulting Center for Statistics at National Cheng Kung University for advice on the statistical analyses. We thank Chih-Ming Hung for the helpful discussions on the data analyses, Yuying Hsu for the early suggestions on the behavioral experimental design, and Yoshihito Hayashi for the valuable advice on the academic writing.

## DECLARATIONS

### Author contributions

Conceptualization, N.O.; methodology, M.-T.C., K.O., A.M., and T.-D.V.; formal analysis, M.-T.C. and K.O.; investigation, M.-T.C. and A.M.; visualization, M.-T.C. writing—original draft, M.-T.C.; writing—review & editing, M.-T.C., T.-D.V., K.O., A.M., H.-V.W., and N.O.; visualization, M.-T.C.; supervision, N.O. and H.-V.W.; project administration, M.-T.C.; resources, H.-V.W. and N.O.; funding acquisition, N.O. and H.-V.W.

### Data and code availability

The RNA-seq datasets generated during this study have been deposited in the DDBJ Sequence Read Archive and are publicly available as of the date of publication under the BioProject accession number PRJDB42482 (sample accession numbers: DRR1061099–DRR1061128). This paper does not report original code; MDC analysis was performed using the published script from Fang et al., with the significance classification criteria and all other computational analyses detailed in the Materials and Methods section using standard, publicly available software packages. Any additional information or raw data required to reanalyze the findings reported in this paper are available from the corresponding author upon reasonable request.

### Declaration of generative AI and AI-assisted technologies in the writing process

During the preparation of this manuscript, the author(s) employed Claude (Anthropic) for rewriting existing text, grammar and phrasing checks, translation support, and discussion of statistical and analytical approaches. This tool served only in an assistive capacity; all original ideas, arguments, data interpretation, and conclusions presented in this work originated with the author(s). All AI-generated content was reviewed, verified, and revised by the author(s), who retain full responsibility for the scientific content, interpretation, and conclusions of this work. No generative AI tool is credited as an author of this work.

## SUPPORTING INFORMATION

Additional supporting information is available in the online version of this article.

**Document S1**. Figures S1–S6 and Tables S1 and S6

**Data S1.** Table S2–S5

## REFERENCES

[1] Hsu Y, Earley RL, Wolf LL. Modulation of aggressive behaviour by fighting experience: mechanisms and contest outcomes. Biological Reviews 2006;81:33–74. 10.1017/S146479310500686X.

[2] Yan JL, Smith NMT, Filice DCS, Dukas R. Winner and loser effects: a meta-analysis. Anim Behav 2024;216:15–22. 10.1016/j.anbehav.2024.07.014.

[3] Fuxjager MJ, Marler CA. How and why the winner effect forms: influences of contest environment and species differences. Behavioral Ecology 2010;21:37–45. 10.1093/beheco/arp148.

[4] Kasumovic MM, Elias DO, Sivalinghem S, Mason AC, Andrade MCB. Examination of prior contest experience and the retention of winner and loser effects. Behavioral Ecology 2010;21:404–9. 10.1093/beheco/arp204.

[5] Lehner SR, Rutte C, Taborsky M. Rats Benefit from Winner and Loser Effects. Ethology 2011;117:949–60. 10.1111/j.1439-0310.2011.01962.x.

[6] Laskowski KL, Wolf M, Bierbach D. The making of winners (And losers): How early dominance interactions determine adult social structure in a clonal fish. Proceedings of the Royal Society B: Biological Sciences 2016;283:20160183. 10.1098/rspb.2016.0183/84446.

[7] Trannoy S, Penn J, Lucey K, Popovic D, Kravitz EA. Short and long-lasting behavioral consequences of agonistic encounters between male *Drosophila melanogaster*. Proc Natl Acad Sci U S A 2016;113:4818–23. 10.1073/pnas.1520953113.

[8] Hock K, Huber R. Models of winner and loser effects: a cost–benefit analysis. Behaviour 2009;146:69–87. 10.1163/156853908X390931.

[9] Rutte C, Taborsky M, Brinkhof MWG. What sets the odds of winning and losing? Trends Ecol Evol 2006;21:16–21. 10.1016/j.tree.2005.10.014.

[10] Zang C, Chung MHJ, Neeman T, Harrison L, Vinogradov IM, Jennions MD. Does losing reduce the tendency to engage with rivals to reach mates? An experimental test. Behavioral Ecology 2024;35:arae037. 10.1093/beheco/arae037.

[11] Huang SP, Yang SY, Hsu Y. Persistence of Winner and Loser Effects Depends on the Behaviour Measured. Ethology 2011;117:171–80. 10.1111/j.1439-0310.2010.01856.x.

[12] Oliveira RF, Silva JF, Simões JM. Fighting zebrafish: Characterization of aggressive behavior and winner-loser effects. Zebrafish 2011;8:73–81. 10.1089/zeb.2011.0690.

[13] Oliveira RF, Silva A, Canário AVM. Why do winners keep winning? Androgen mediation of winner but not loser effects in cichlid fish. Proc Biol Sci 2009;276:2249–56. 10.1098/rspb.2009.0132.

[14] Filby AL, Paull GC, Hickmore TFA, Tyler CR. Unravelling the neurophysiological basis of aggression in a fish model. BMC Genomics 2010;11:498. 10.1186/1471-2164-11-498.

[15] Sneddon LU, Schmidt R, Fang Y, Cossins AR. Molecular Correlates of Social Dominance: A Novel Role for Ependymin in Aggression. PLoS One 2011;6:e18181. 10.1371/journal.pone.0018181.

[16] Oliveira RF, Simões JM, Teles MC, Oliveira CR, Becker JD, Lopes JS. Assessment of fight outcome is needed to activate socially driven transcriptional changes in the zebrafish brain. Proceedings of the National Academy of Sciences 2016;113:E654–61. 10.1073/pnas.1514292113.

[17] Renn SCP, Aubin-Horth N, Hofmann HA. Fish and chips: functional genomics of social plasticity in an African cichlid fish. Journal of Experimental Biology 2008;211:3041–56. 10.1242/jeb.018242.

[18] de la Fuente A. From ‘differential expression’ to ‘differential networking’ – identification of dysfunctional regulatory networks in diseases. Trends in Genetics 2010;26:326–33. 10.1016/j.tig.2010.05.001.

[19] Zhang B, Gaiteri C, Bodea LG, Wang Z, McElwee J, Podtelezhnikov AA, et al. Integrated systems approach identifies genetic nodes and networks in late-onset Alzheimer’s disease. Cell 2013;153:707–20. 10.1016/j.cell.2013.03.030.

[20] Rittschof CC, Bukhari SA, Sloofman LG, Troy JM, Caetano-Anollés D, Cash-Ahmed A, et al. Neuromolecular responses to social challenge: Common mechanisms across mouse, stickleback fish, and honey bee. Proceedings of the National Academy of Sciences 2014;111:17929–34. 10.1073/pnas.1420369111.

[21] Adeoye T, Shah SI, Ullah G. Systematic Analysis of Biological Processes Reveals Gene Co-expression Modules Driving Pathway Dysregulation in Alzheimer’s Disease. Aging Dis 2024;16:1598. 10.14336/ad.2024.0429.

[22] Fang YT, Kuo HC, Chen CY, Chou SJ, Lu CW, Hung CM. Brain Gene Regulatory Networks Coordinate Nest Construction in Birds. Mol Biol Evol 2024;41:msae125. 10.1093/molbev/msae125.

[23] Robinson GE. Integrative animal behaviour and sociogenomics. Trends Ecol Evol 1999;14:202–5. 10.1016/S0169-5347(98)01536-5.

[24] Robinson GE, Grozinger CM, Whitfield CW. Sociogenomics: Social life in molecular terms. Nat Rev Genet 2005;6:257–70. 10.1038/nrg1575.

[25] Robinson GE, Fernald RD, Clayton DF. Genes and social behavior. Science (1979) 2008;322:896–900. 10.1126/science.1159277.

[26] Chandrasekaran S, Ament SA, Eddy JA, Rodriguez-Zas SL, Schatz BR, Price ND, et al. Behavior-specific changes in transcriptional modules lead to distinct and predictable neurogenomic states. Proc Natl Acad Sci U S A 2011;108:18020–5. 10.1073/pnas.1114093108.

[27] Shpigler HY, Saul MC, Murdoch EE, Cash-Ahmed AC, Seward CH, Sloofman L, et al. Behavioral, transcriptomic and epigenetic responses to social challenge in honey bees. Genes Brain Behav 2017;16:579–91. 10.1111/gbb.12379.

[28] Hamilton AR, Traniello IM, Ray AM, Caldwell AS, Wickline SA, Robinson GE. Division of labor in honey bees is associated with transcriptional regulatory plasticity in the brain. Journal of Experimental Biology 2019;222:jeb200196. 10.1242/jeb.200196/20799.

[29] Sinha S, Jones BM, Traniello IM, Bukhari SA, Halfon MS, Hofmann HA, et al. Behavior-related gene regulatory networks: A new level of organization in the brain. Proc Natl Acad Sci U S A 2020;117:23270–9. 10.1073/pnas.1921625117.

[30] Jones BM, Rao VD, Gernat T, Jagla T, Cash-Ahmed AC, Rubin BER, et al. Individual differences in honey bee behavior enabled by plasticity in brain gene regulatory networks. Elife 2020;9:1–28. 10.7554/elife.62850.

[31] Regan CT. The Asiatic fishes of the family Anabantidae. Proceedings of the Zoological Society of London 1910;1909:767–87.

[32] Simpson MJA. The Display of the Siamese Fighting Fish, Betta splendens. Animal Behaviour Monographs 1968;1:i–73. 10.1016/S0066-1856(68)80001-9.

[33] Bronstein PM. Onset of combat in male *Betta splendens*. J Comp Psychol 1983;97:135–9. 10.1037/0735-7036.97.2.135.

[34] Karino K, Someya C. The influence of sex, line, and fight experience on aggressiveness of the Siamese fighting fish in intrasexual competition. Behavioural Processes 2007;75:283–9. 10.1016/j.beproc.2007.03.002.

[35] Fan G, Chan J, Ma K, Yang B, Zhang H, Yang X, et al. Chromosome-level reference genome of the Siamese fighting fish *Betta splendens*, a model species for the study of aggression. Gigascience 2018;7:1–7. 10.1093/gigascience/giy087.

[36] Srikulnath K, Singchat W, Laopichienpong N, Ahmad SF, Jehangir M, Subpayakom N, et al. Overview of the betta fish genome regarding species radiation, parental care, behavioral aggression, and pigmentation model relevant to humans. Genes & Genomics 2021 43:2 2021;43:91–104. 10.1007/S13258-020-01027-2.

[37] Kwon YM, Vranken N, Hoge C, Lichak MR, Norovich AL, Francis KX, et al. Genomic consequences of domestication of the Siamese fighting fish. Sci Adv 2022;8:4950. 10.1126/sciadv.abm4950.

[38] Vu T-D, Iwasaki Y, Oshima K, Chiu M-T, Nikaido M, Okada N. A unique neurogenomic state emerges after aggressive confrontations in males of the fish *Betta splendens*. Gene 2021;784:145601. 10.1016/j.gene.2021.145601.

[39] Vu T-D, Iwasaki Y, Shigenobu S, Maruko A, Oshima K, Iioka E, et al. Behavioral and brain-transcriptomic synchronization between the two opponents of a fighting pair of the fish *Betta splendens*. PLoS Genet 2020;16:e1008831-.

[40] Vu T-D, Oshima K, Matsumura K, Iwasaki Y, Chiu M-T, Nikaido M, et al. Alternative splicing plays key roles in response to stress across different stages of fighting in the fish *Betta splendens*. BMC Genomics 2022;22:920. 10.1186/s12864-022-08609-2.

[41] Earley RL, Hsu Y, Wolf LL. The Use of Standard Aggression Testing Methods to Predict Combat Behaviour and Contest Outcome in *Rivulus marmoratus* Dyads (Teleostei: Cyprinodontidae). Ethology 2000;106:743–61. 10.1046/j.1439-0310.2000.00586.x.

[42] Hsu Y, Wolf LL. The winner and loser effect: Integrating multiple experiences. Anim Behav 1999;57:903–10. 10.1006/anbe.1998.1049.

[43] Earley RL, Lu CK, Lee IH, Wong SC, Hsu Y. Winner and loser effects are modulated by hormonal states. Front Zool 2013;10:6. 10.1186/1742-9994-10-6.

[44] Hsu Y, Wolf LL. The winner and loser effect: what fighting behaviours are influenced? Anim Behav 2001;61:777–86. 10.1006/anbe.2000.1650.

[45] Okada K, Okada Y, Dall SRX, Hosken DJ. Loser-effect duration evolves independently of fighting ability. Proc Biol Sci 2019;286:20190582. 10.1098/rspb.2019.0582.

[46] Hubená P, Benrejdal L, Brodin D, Axling J, Sarma O Sen, Bergman P, et al. Effects of Stress Coping Styles and Social Defeat on Zebrafish Behaviour and Brain Transcriptomics. Neurosci Bull 2025;42:989. 10.1007/S12264-025-01506-0.

[47] Bagot RCC, Cates HMM, Purushothaman I, Lorsch ZSS, Walker DMM, Wang J, et al. Circuit-wide Transcriptional Profiling Reveals Brain Region-Specific Gene Networks Regulating Depression Susceptibility. Neuron 2016;90:969–83. 10.1016/j.neuron.2016.04.015.

[48] Winberg S, Nilsson GE. Roles of brain monoamine neurotransmitters in agonistic behaviour and stress reactions, with particular reference to fish. Comp Biochem Physiol C Pharmacol Toxicol Endocrinol 1993;106:597–614. 10.1016/0742-8413(93)90216-8.

[49] Øvrli Ø, Harris CA, Winberg S. Short-Term Effects of Fights for Social Dominance and the Establishment of Dominant-Subordinate Relationships on Brain Monoamines and Cortisol in Rainbow Trout. Brain Behav Evol 1999;54:263–75. 10.1159/000006627.

[50] Saul MC, Seward CH, Troy JM, Zhang H, Sloofman LG, Lu X, et al. Transcriptional regulatory dynamics drive coordinated metabolic and neural response to social challenge in mice. Genome Res 2017;27:959–72. 10.1101/gr.214221.116.

[51] Uy FMK, Jernigan CM, Zaba NC, Mehrotra E, Miller SE, Sheehan MJ. Dynamic neurogenomic responses to social interactions and dominance outcomes in female paper wasps. PLoS Genet 2021;17:e1009474. 10.1371/journal.pgen.1009474.

[52] Quackenbush J. Computational analysis of microarray data. Nature Reviews Genetics 2001 2:6 2001;2:418–27. 10.1038/35076576.

[53] Fuxjager MJ, Forbes-Lorman RM, Coss DJ, Auger CJ, Auger AP, Marler CA. Winning territorial disputes selectively enhances androgen sensitivity in neural pathways related to motivation and social aggression. Proc Natl Acad Sci U S A 2010;107:12393–8. 10.1073/pnas.1001394107.

[54] Lindberg D, Shan D, Ayers-Ringler J, Oliveros A, Benitez J, Prieto M, et al. Purinergic Signaling and Energy Homeostasis in Psychiatric Disorders. Curr Mol Med 2015;15:275. 10.2174/1566524015666150330163724.

[55] Dai B, Zheng B, Dai X, Cui X, Yin L, Cai J, et al. Experience-dependent dopamine modulation of male aggression. Nature 2025 639:8054 2025;639:430–7. 10.1038/s41586-024-08459-w.

[56] Malenka RC, Bear MF. LTP and LTD: An embarrassment of riches. Neuron 2004;44:5–21. 10.1016/j.neuron.2004.09.012.

[57] Huganir RL, Nicoll RA. AMPARs and Synaptic Plasticity: The Last 25 Years. Neuron 2013;80:704–17. 10.1016/j.neuron.2013.10.025.

[58] Backström T, Winberg S. Serotonin coordinates responses to social stress-What we can learn from fish. Front Neurosci 2017;11:294719. 10.3389/fnins.2017.00595.

[59] Zhou T, Zhu H, Fan Z, Wang F, Chen Y, Liang H, et al. History of winning remodels thalamo-PFC circuit to reinforce social dominance. Science (1979) 2017;357:162–8. 10.1126/science.aak9726.

[60] Chou MY, Amo R, Kinoshita M, Cherng BW, Shimazaki H, Agetsuma M, et al. Social conflict resolution regulated by two dorsal habenular subregions in zebrafish. Science (1979) 2016;352:87–90. 10.1126/science.aac9508.

[61] Guo H, Dixon B. Understanding acute stress-mediated immunity in teleost fish. Fish and Shellfish Immunology Reports 2021;2:100010. 10.1016/j.fsirep.2021.100010.

[62] Takahashi A. Associations of the immune system in aggression traits and the role of microglia as mediators. Neuropharmacology 2024;256:110021. 10.1016/j.neuropharm.2024.110021.

[63] Azeredo R, Machado M, Pereiro P, Barany A, Mancera JM, Costas B. Acute Inflammation Induces Neuroendocrine and Opioid Receptor Genes Responses in the Seabass *Dicentrarchus labrax* Brain. Biology (Basel) 2022;11:364. 10.3390/biology11030364/S1.

[64] Shimo Y, Cathomas F, Lin HY, Chan KL, Parise LF, Li L, et al. Social stress induces autoimmune responses against the brain. Proc Natl Acad Sci U S A 2023;120:e2305778120. 10.1073/pnas.2305778120.

[65] Yang J, Jia Y, Guo T, Zhang S, Huang J, Lu H, et al. Comparative Analysis of HPA-Axis Dysregulation and Dynamic Molecular Mechanisms in Acute Versus Chronic Social Defeat Stress. Int J Mol Sci 2025;26:6063. 10.3390/ijms26136063.

[66] Archie EA, Altmann J, Alberts SC. Social status predicts wound healing in wild baboons. Proc Natl Acad Sci U S A 2012;109:9017–22. 10.1073/pnas.1206391109.

[67] Takahashi A, Flanigan ME, McEwen BS, Russo SJ. Aggression, social stress, and the immune system in humans and animal models. Front Behav Neurosci 2018;12:338283. 10.3389/fnbeh.2018.00056.

[68] Leschak CJ, Eisenberger NI. Two Distinct Immune Pathways Linking Social Relationships With Health: Inflammatory and Antiviral Processes. Psychosom Med 2019;81:711–9. 10.1097/psy.0000000000000685.

[69] Stevenson PA, Rillich J. Isolation Associated Aggression – A Consequence of Recovery from Defeat in a Territorial Animal. PLoS One 2013;8:e74965. 10.1371/journal.pone.0074965.

[70] De Groot J, Boersma WJA, Scholten JW, Koolhaas JM. Social stress in male mice impairs long-term antiviral immunity selectively in wounded subjects. Physiol Behav 2002;75:277– 85. 10.1016/S0031-9384(01)00677-1.

[71] De Groot J, Van Milligen FJ, Moonen-Leusen BWM, Thomas G, Koolhaas JM. A single social defeat transiently suppresses the anti-viral immune response in mice. J Neuroimmunol 1999;95:143–51. 10.1016/S0165-5728(99)00005-3.

[72] Price J, Sloman L, Gardner R, Gilbert P, Rohde P. The Social Competition Hypothesis of Depression. British Journal of Psychiatry 1994;164:309–15. 10.1192/bjp.164.3.309.

[73] Stewart A, Kadri F, DiLeo J, Min Chung K, Cachat J, Goodspeed J, et al. The Developing Utility of Zebrafish in Modeling Neurobehavioral Disorders. Int J Comp Psychol 2010;23. 10.46867/ijcp.2010.23.01.01.

[74] Balzarini V, Taborsky M, Wanner S, Koch F, Frommen JG. Mirror, mirror on the wall: the predictive value of mirror tests for measuring aggression in fish. Behavioral Ecology and Sociobiology 2014 68:5 2014;68:871–8. 10.1007/S00265-014-1698-7.

[75] Friard O, Gamba M. BORIS: a free, versatile open-source event-logging software for video/audio coding and live observations. Methods Ecol Evol 2016;7:1325–30. 10.1111/2041-210X.12584.

[76] Hofmann HA, Schildberger K. Assessment of strength and willingness to fight during aggressive encounters in crickets. Anim Behav 2001;62:337–48. 10.1006/anbe.2001.1746.

[77] Matthews M, Varga ZM. Anesthesia and Euthanasia in Zebrafish. ILAR J 2012;53:192–204. 10.1093/ilar.53.2.192.

[78] Martin M. Cutadapt removes adapter sequences from high-throughput sequencing reads. EMBnet J 2011;17:10–2. 10.14806/EJ.17.1.200.

[79] Bolger AM, Lohse M, Usadel B. Trimmomatic: a flexible trimmer for Illumina sequence data. Bioinformatics 2014;30:2114. 10.1093/bioinformatics/btu170.

[80] Langmead B, Salzberg SL. Fast gapped-read alignment with Bowtie 2. Nat Methods 2012;9:357. 10.1038/nmeth.1923.

[81] Kim D, Paggi JM, Park C, Bennett C, Salzberg SL. Graph-based genome alignment and genotyping with HISAT2 and HISAT-genotype. Nature Biotechnology 2019 37:8 2019;37:907–15. 10.1038/s41587-019-0201-4.

[82] Li H, Handsaker B, Wysoker A, Fennell T, Ruan J, Homer N, et al. The Sequence Alignment/Map format and SAMtools. Bioinformatics 2009;25:2078. 10.1093/bioinformatics/btp352.

[83] Liao Y, Smyth GK, Shi W. featureCounts: an efficient general purpose program for assigning sequence reads to genomic features. Bioinformatics 2014;30:923–30. 10.1093/bioinformatics/btt656.

[84] Robinson MD, McCarthy DJ, Smyth GK. edgeR: a Bioconductor package for differential expression analysis of digital gene expression data. Bioinformatics 2009;26:139. 10.1093/bioinformatics/btp616.

[85] Suzuki R, Shimodaira H. Pvclust: an R package for assessing the uncertainty in hierarchical clustering. Bioinformatics 2006;22:1540–2. 10.1093/bioinformatics/btl117.

[86] Gu Z, Eils R, Schlesner M. Complex heatmaps reveal patterns and correlations in multidimensional genomic data. Bioinformatics 2016;32:2847–9. 10.1093/bioinformatics/btw313.

[87] Langfelder P, Horvath S. WGCNA: an R package for weighted correlation network analysis. BMC Bioinformatics 2008 9:1 2008;9:559-. 10.1186/1471-2105-9-559.

[88] Chen Y, Lun ATL, Smyth GK. From reads to genes to pathways: differential expression analysis of RNA-Seq experiments using Rsubread and the edgeR quasi-likelihood pipeline. F1000Res 2016;5:1438. 10.12688/f1000reasearch.8987.2.

[89] Hsu Y, Earley RL, Wolf LL. Aggressive Behaviour in Fish: Integrating Information about Contest Costs. Fish Cognition and Behavior 2011:108–34. 10.1002/9781444342536.CH6.

[90] Sherman BT, Hao M, Qiu J, Jiao X, Baseler MW, Lane HC, et al. DAVID: a web server for functional enrichment analysis and functional annotation of gene lists (2021 update). Nucleic Acids Res 2022;50:W216–21. 10.1093/nar/gkac194.

[91] Huang DW, Sherman BT, Lempicki RA. Systematic and integrative analysis of large gene lists using DAVID bioinformatics resources. Nat Protoc 2009;4:44–57. 10.1038/nprot.2008.211.

[92] Kuznetsova A, Brockhoff PB, Christensen RHB. lmerTest Package: Tests in Linear Mixed Effects Models. J Stat Softw 2017;82:1–26. 10.18637/jss.v082.I13.

[93] Favati A, Løvlie H, Leimar O. Individual aggression, but not winner–loser effects, predicts social rank in male domestic fowl. Behavioral Ecology 2017;28:874–82. 10.1093/beheco/arx053.

[94] Lenth R V., Piaskowski J. emmeans: Estimated Marginal Means, aka Least-Squares Means. 2026; R package version 2.0.3.

