## Supplementary material for "Asymmetric Neurogenomic States Emerge in Winners and Losers After Social Competition": Document S1

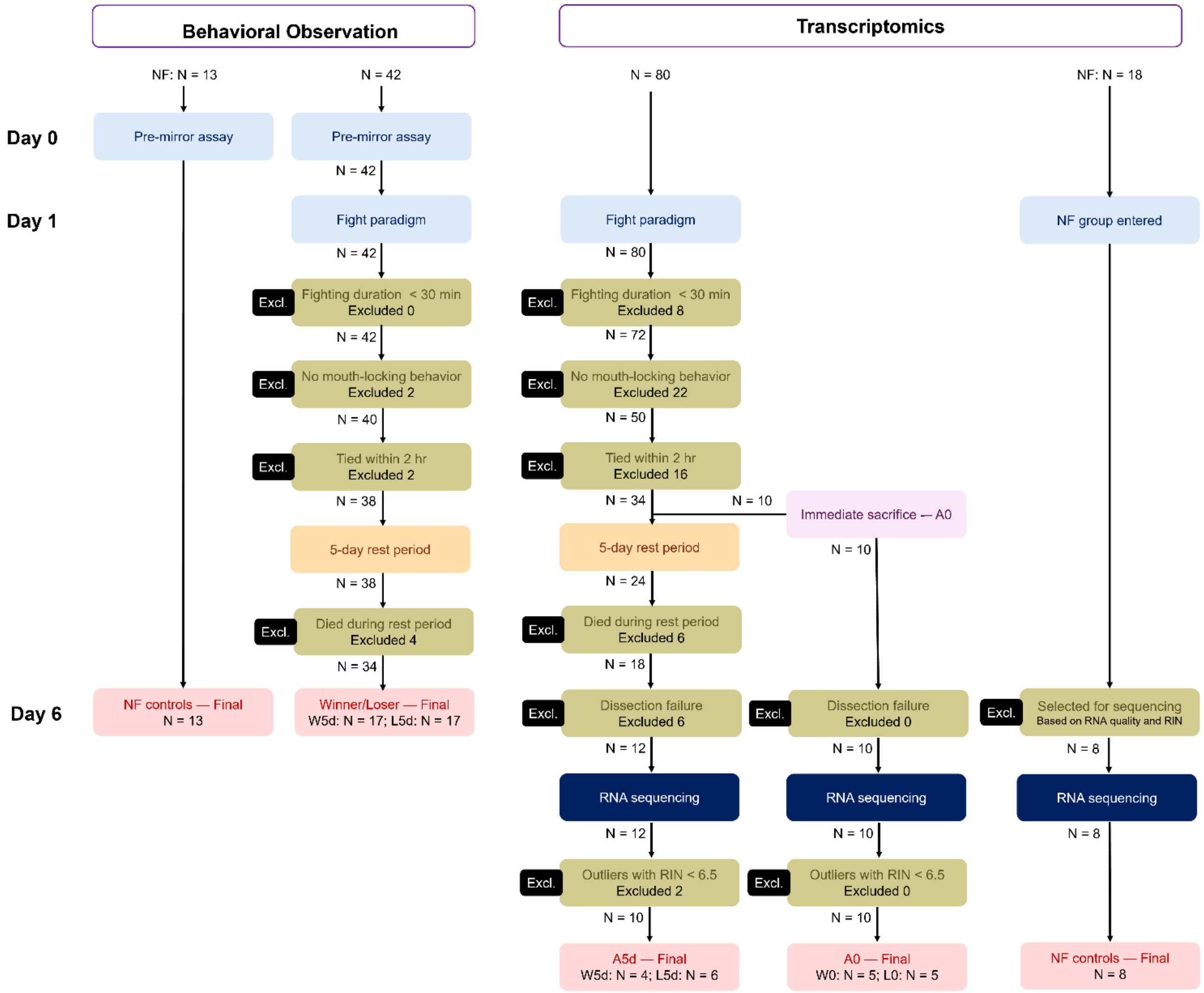


**Figure S1. Sample attrition and inclusion criteria for behavioral and transcriptomic analyses.** Flow diagram showing sample sizes at each stage of screening for behavioral observation and transcriptomic analyses, each comprising fighting and nonfighting control (NF) groups. Fighting fish were sequentially screened for fighting duration (≥30 min), presence of mouth-locking (the highest-intensity aggressive behavior, used to standardize fighting experience), and resolution of the contest within a 2-h observation period (unresolved pairs were excluded as ties). For transcriptomics, a subset of fish was sacrificed immediately after fighting (A0), while the remainder underwent a 5-day rest period before sampling (A5d); when one individual of a pair died during the remaining period, its opponent was also excluded to maintain paired winner-loser sampling. Dissection failure refers to unsuccessful whole-brain extraction or inadequate tissue quality. Following RNA sequencing, samples with an RNA integrity number (RIN) < 6.5 were excluded as outliers where sample size allowed. The final sample sizes correspond to those reported in Table S1.

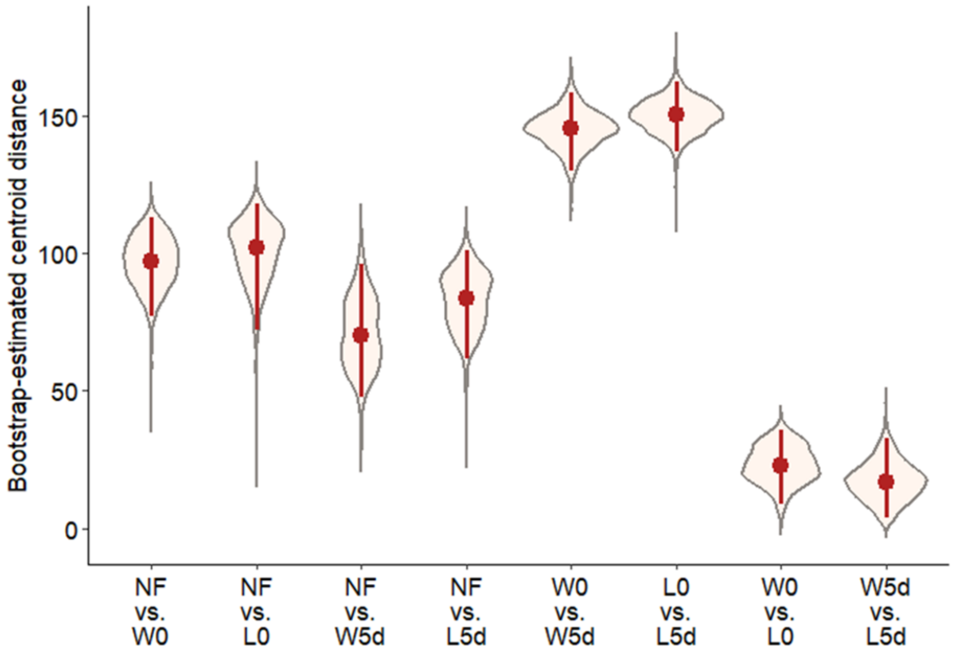


**Figure S2. Bootstrap-estimated centroid distances between group pairs in PCA space**

Violin plots show the distribution of bootstrap-estimated Euclidean centroid distances between each pair of groups across 1,000 bootstrap resampling iterations in the PC1–PC2 space. Red dots indicate the median distance, and red lines indicate the interquartile range. Group pairs are organized into three categories: NF versus postfight groups (NF vs. W0, NF vs. L0, NF vs. W5d, and NF vs. L5d), between-stage comparisons within the same outcome (W0 vs. W5d and L0 vs. L5d), and within-stage winner–loser comparisons (W0 vs. L0 and W5d vs. L5d).

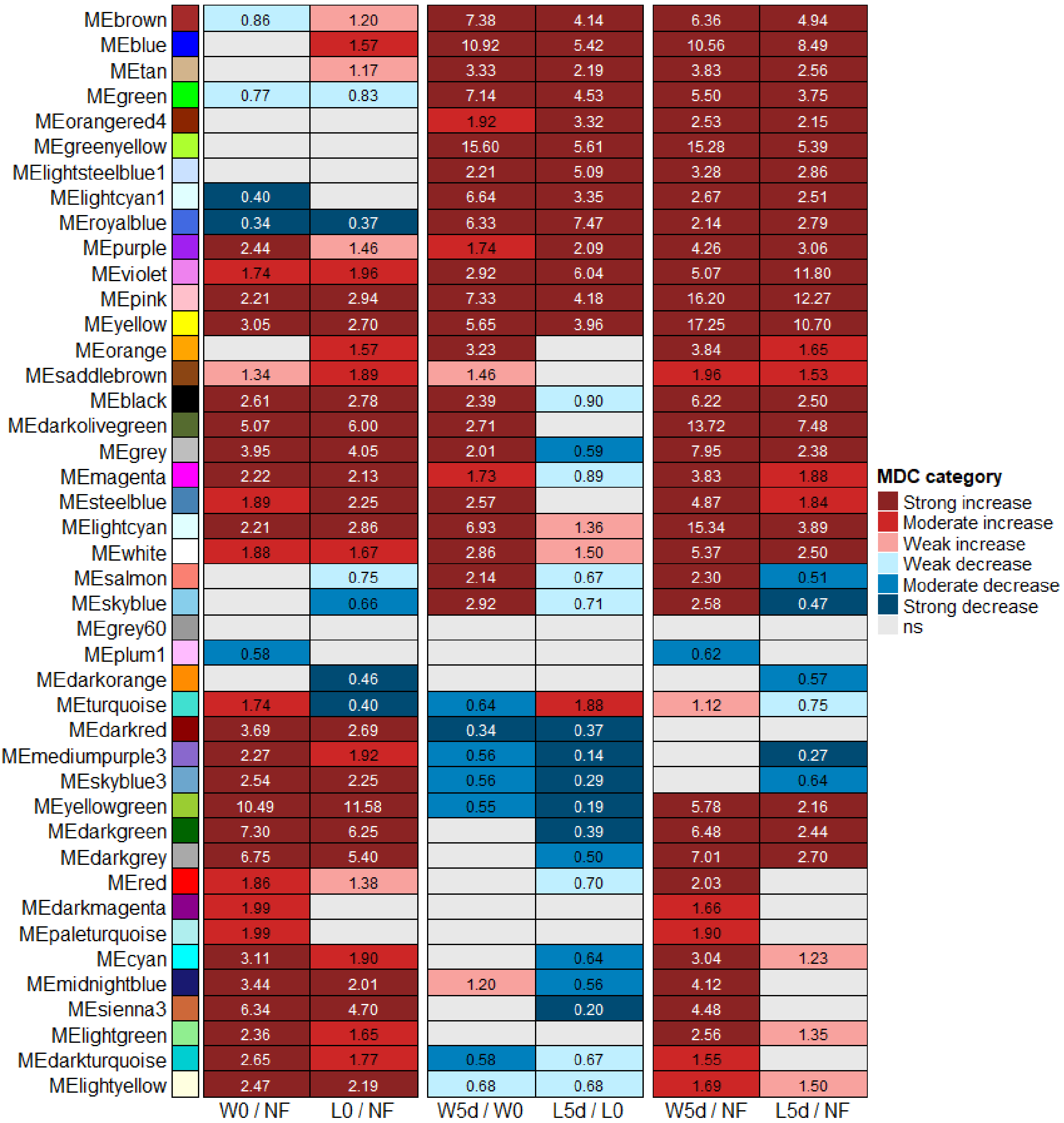


**Figure S3. Modular differential connectivity (MDC) across all 43 WGCNA modules.** Heatmap showing graded MDC classification (strong, moderate, or weak increase or decrease in connectivity, or not significant [ns]) for all 43 coexpression modules across three sets of pairwise comparisons, corresponding to Figure 3: immediate postfight versus controls (W0/NF and L0/NF), temporal transitions from immediate to later postfight (W5d/W0 and L5d/L0), and later postfight versus controls (W5d/NF and L5d/NF). The values in the cells indicate the MDC values, shown only for significant comparisons (FDR < 0.05). The 20 modules retained for the main analysis (Figure 3) were selected after excluding modules with significant batch effects on eigengene expression (FDR < 0.05; Methods).

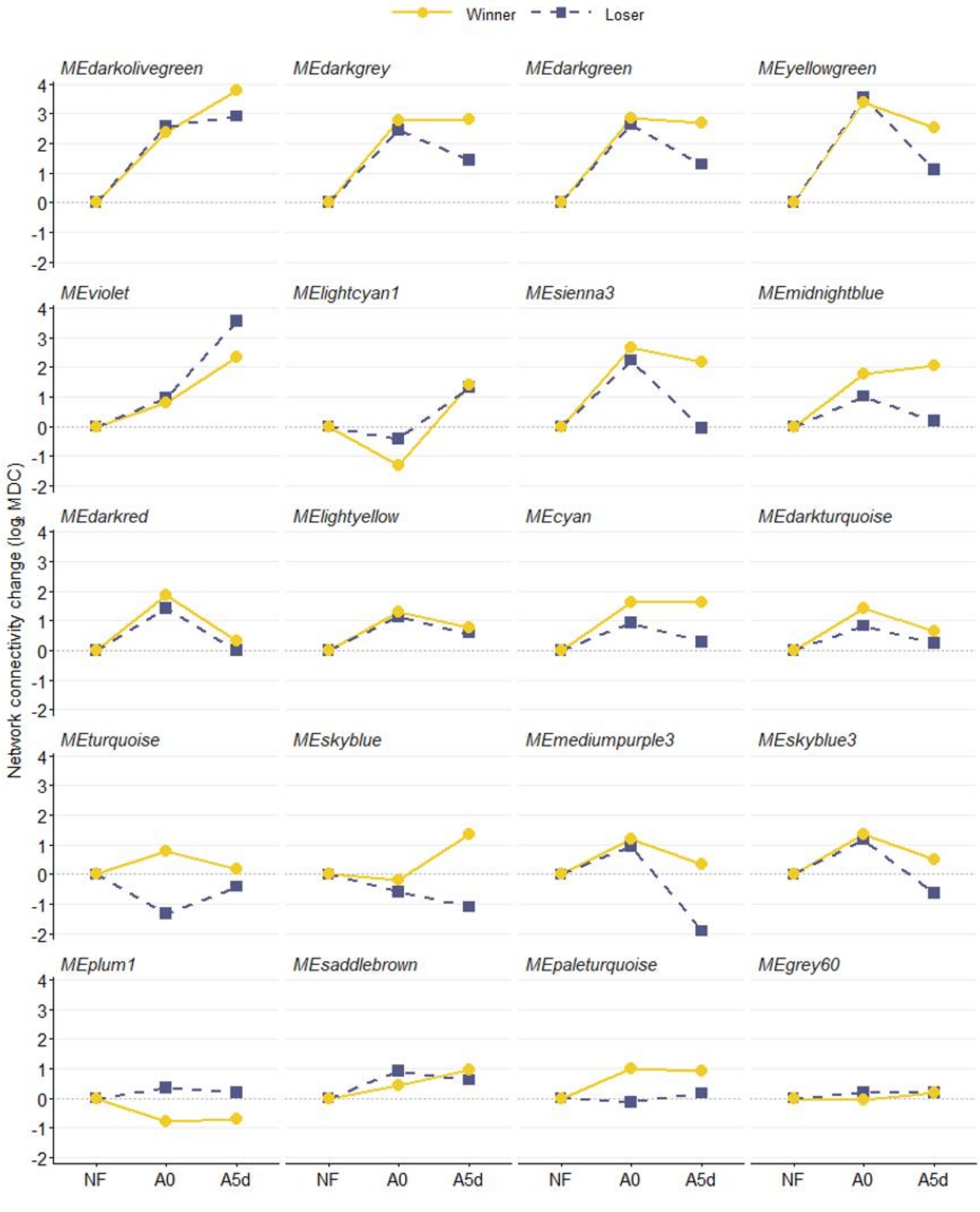


**Figure S4. Network connectivity trajectories across all 20 retained modules.**
Network connectivity change ($log_2$MDC, relative to NF) across the NF, immediate postfight (A0), and later postfight (A5d) states for winners (yellow, solid) and losers (dark blue, dashed) in all 20 modules retained for the main analysis (Figure 3), including the four representative modules shown in Figure 3B–E. NF was set to a reference value of 0 (MDC = 1); the A0 and A5d values represent the $log_2$MDC of winners or losers relative to that of NF at the corresponding time points (i.e., W0/NF and L0/NF for A0; W5d/NF and L5d/NF for A5d).

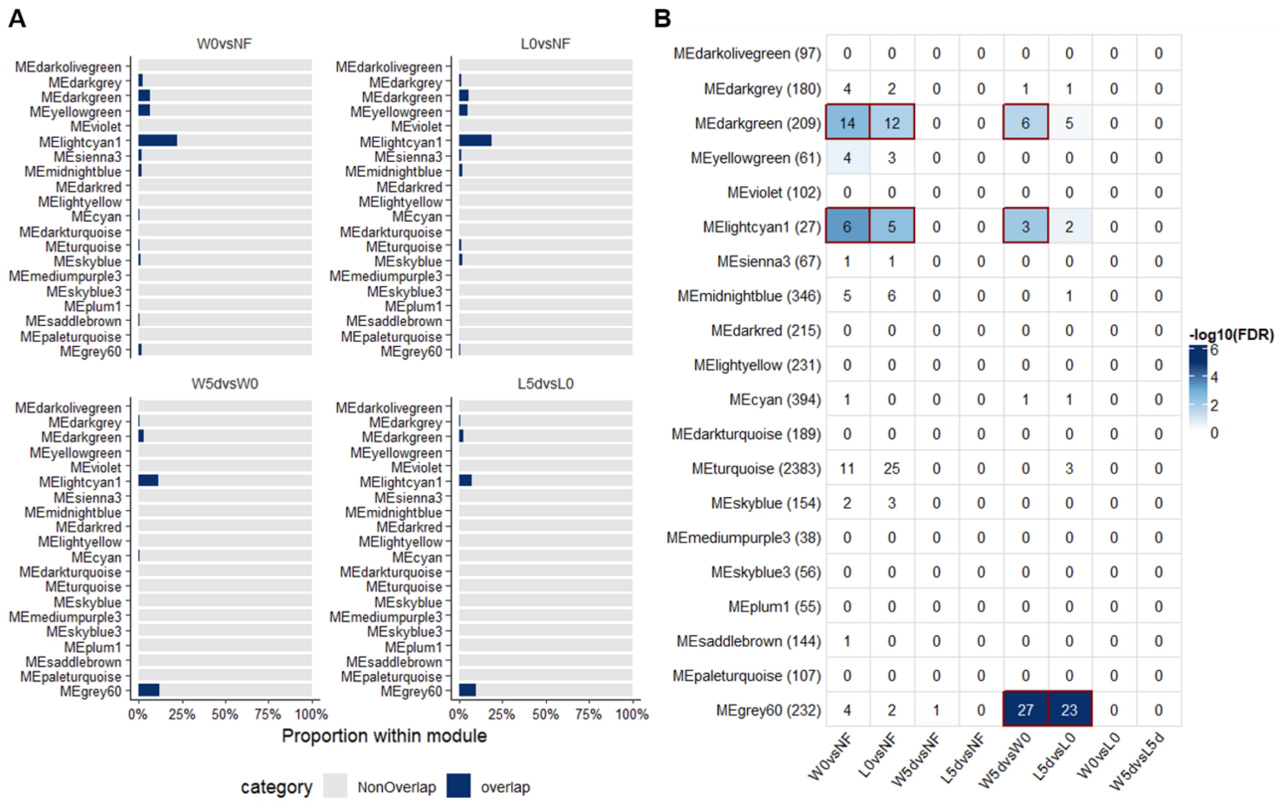


**Figure S5. Differential gene expression does not account for network connectivity changes.** (A) Proportion of module genes overlapping with DEGs (up- and downregulated combined) for the acute (W0 vs. NF, L0 vs. NF) and transition (W5d vs. W0, L5d vs. L0) stage comparisons, across the 20 modules retained in Figure 3A (module order as in Figure 3A). Overlap (navy) and nonoverlap (gray) proportions are shown relative to the total number of genes in each module. (B) Hypergeometric enrichment of DEGs (up- and downregulated combined; background = 22,296 filtered genes) within each module across all pairwise comparisons. The cell values indicate the number of overlapping genes; the color indicates $-\log_{10}$FDR (Benjamini–Hochberg correction applied separately for each comparison); and the red outlines indicate an FDR < 0.05. The row labels indicate module identity and the total number of genes per module. The numbers of DEGs used for each comparison are detailed in Figure 2D.

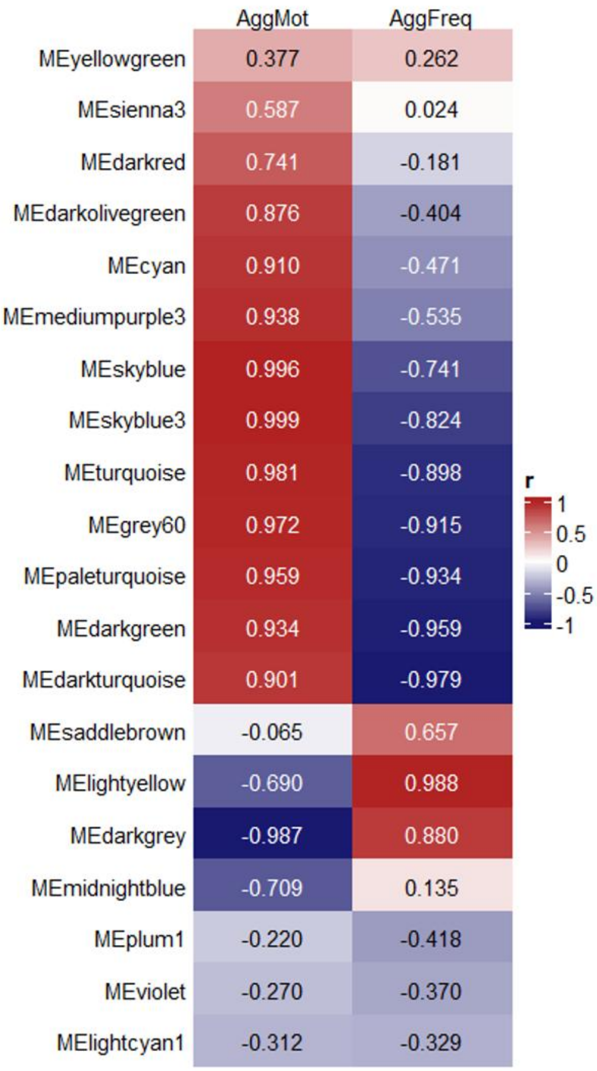


**Figure S6. Module eigengene correlations with aggressive behavior.**

Heatmap of Pearson correlation coefficients (*r*) between module eigengenes (the first principal component of module gene expression) and aggressive motivation (*AggMot*, *-AggLat*) or aggressive output (*AggFreq*) based on group-mean values for the NF, winner (W5d), and loser (L5d) groups (n = 3 group-level points per module; exploratory given the limited number of group-level observations). Red indicates a positive correlation, and blue indicates a negative correlation; the values in the cells indicate the *r* values. This analysis provides the complete set of correlation coefficients underlying the binarized behavior annotations ($|r| > 0.8$) shown in Figure 3A.

**Table S1. Sample numbers used in the behavioral and transcriptomic analyses**

| | Behavioral data | RNA-seq |
|---|---|---|
| Immediate postfight (A0) | | |
| - Immediate winners (W0) | - | 5 |
| - Immediate losers (L0) | - | 5 |
| Later postfight (A5d) | | |
| - Winner effect (W5d) | 17 | 4 |
| - Loser effect (L5d) | 17 | 6 |
| Nonfighting controls (NF) | 13 | 8 |

*A0 samples did not undergo mirror assays, as prior experience indicates that fish are physically exhausted immediately after fighting and show little to no response to mirror stimuli.*
*Full sample attrition details are provided in Fig. S1.*

**Table S6. WGCNA soft-thresholding power sensitivity analysis: Jaccard similarity of module gene membership relative to power = 8**

| Module (power = 8) | Size | Jaccard | | |
|---|---|---|---|---|
| | | vs. power = 6 | vs. power = 9 | vs. power = 12 |
| **MEturquoise** | **2,383** | **0.965** | **0.829** | **0.777** |
| MEblue | 1,887 | 0.862 | 0.775 | 0.760 |
| MEbrown | 1,854 | 0.430 | 0.807 | 0.802 |
| MEyellow | 1,718 | 0.736 | 0.796 | 0.720 |
| MEgreen | 1,332 | 0.862 | 0.541 | 0.428 |
| MEred | 1,190 | 0.861 | 0.879 | 0.832 |
| MEblack | 1,063 | 0.804 | 0.924 | 0.707 |
| MEpink | 1,045 | 0.379 | 0.880 | 0.451 |
| MEmagenta | 911 | 0.757 | 0.644 | 0.672 |
| MEpurple | 656 | 0.852 | 0.900 | 0.538 |
| MEgreenyellow | 563 | 0.872 | 0.957 | 0.807 |
| MEtan | 498 | 0.340 | 0.191 | 0.548 |
| MEsalmon | 469 | 0.832 | 0.892 | 0.752 |
| **MEcyan** | **394** | **0.608** | **0.573** | **0.377** |
| **MEmidnightblue** | **346** | **0.781** | **0.689** | **0.658** |
| MElightcyan | 327 | 0.465 | 0.659 | 0.642 |
| **MEgrey60** | **232** | **0.700** | **0.595** | **0.674** |
| MElightgreen | 231 | 0.558 | 0.757 | 0.655 |
| **MElightyellow** | **231** | **0.855** | **0.875** | **0.658** |
| MEroyalblue | 218 | 0.800 | 0.916 | 0.801 |
| **MEdarkred** | **215** | **0.459** | **0.562** | **0.516** |
| **MEdarkgreen** | **209** | **0.681** | **0.811** | **0.399** |
| **MEdarkturquoise** | **189** | **0.813** | **0.865** | **0.738** |
| **MEdarkgrey** | **180** | **0.621** | **0.661** | **0.579** |
| MEorange | 174 | 0.812 | 0.789 | 0.588 |
| MEdarkorange | 166 | 0.785 | 0.738 | 0.753 |
| MEwhite | 155 | 0.060 | 0.154 | 0.469 |
| **MEskyblue** | **154** | **0.034** | **0.576** | **0.500** |
| **MEsaddlebrown** | **144** | **0.604** | **0.693** | **0.569** |
| MEsteelblue | 132 | 0.288 | 0.644 | 0.393 |
| **MEpaleturquoise** | **107** | **0.425** | **0.604** | **0.545** |
| **MEviolet** | **102** | **0.837** | **0.916** | **0.845** |
| **MEdarkolivegreen** | **97** | **0.253** | **0.782** | **0.664** |
| MEdarkmagenta | 75 | 0.578 | 0.413 | 0.495 |
| **MEsienna3** | **67** | **0.710** | **0.844** | **0.725** |
| **MEyellowgreen** | **61** | **0.562** | **0.712** | **0.048** |
| **MEskyblue3** | **56** | **0.402** | **0.542** | **0.418** |
| **MEplum1** | **55** | **0.004** | **0.703** | **0.536** |
| MEorangered4 | 38 | 0.028 | 0.589 | 0.625 |
| **MEmediumpurple3** | **38** | **0.769** | **0.850** | **0.804** |

| Module (power = 8) | Size | Jaccard | | |
|---|---|---|---|---|
| | | vs. power = 6 | vs. power = 9 | vs. power = 12 |
| MElightsteelblue1 | 32 | 0.149 | 0.147 | 0.488 |
| **MElightcyan1** | **27** | **0.750** | **0.771** | **0.675** |

*Jaccard similarity was computed between the gene membership of each module identified at power = 8 and its best-matching module at powers 6, 9, and 12, using the formula* $|A \cap B|/|A \cup B|$*, where A and B denote the gene sets of the two matched modules. The best-matching modules were identified by maximizing gene membership overlap. The modules are sorted by size (number of genes) in descending order. The modules retained for downstream MDC analysis are indicated in bold. The cells with Jaccard similarity < 0.5 are shaded gray, indicating that fewer than half of the genes were shared between matched modules across power settings. Jaccard similarity of 0 or near 0 indicates that the corresponding module was not recoverable at that power setting.*
